# HDAC1 controls the generation and maintenance of effector-like CD8^+^ T cells during chronic viral infection

**DOI:** 10.1101/2024.02.28.580886

**Authors:** Ramona Rica, Monika Waldherr, Marlene Schülein, Emi Miyakoda, Lisa Sandner, Valentina Stolz, Darina Waltenberger, Thomas Krausgruber, Christoph Bock, Nicole Boucheron, Wilfried Ellmeier, Shinya Sakaguchi

## Abstract

CD8^+^ T cell exhaustion is a complex process that involves the differentiation of persistently activated CD8^+^ T cells into functionally distinct cell subsets. Here, we investigated the role of the key epigenetic regulator histone deacetylase 1 (HDAC1) in the differentiation of exhausted T (Tex) cells during chronic viral infection. We uncovered that HDAC1 controls the generation and maintenance of effector-like CX3CR1^+^ Tex cells in a CD8^+^ T cell-intrinsic manner. Deletion of HDAC1 led to expansion of an alternative Tex cell subset characterized by high expression of T cell exhaustion markers, and this was accompanied by elevated viremia. HDAC1 knockout altered the chromatin landscape in progenitor Tex cells, abrogated the expression of effector-like signature genes and interfered with cell fate specification toward the CX3CR1^+^ Tex cell subset. We conclude that HDAC1 is functionally required for controlling viral load during chronic infection by ensuring adequate CX3CR1^+^ Tex cell subset differentiation.

**Highlights:**

- HDAC1 promotes the generation of CX3CR1^+^ effector-like Tex cell subsets in chronic viral infection in a CD8^+^ T cell-intrinsic manner.
- Deletion of HDAC1 leads to an increase of a cell subset enriched in exhaustion markers and is accompanied with elevated viremia.
- HDAC1 is required for the maintenance of the CX3CR1^+^ Tex cell pool.
- HDAC1 deletion alters the chromatin landscape at effector-like signature gene loci in progenitor Tex cells.

## Introduction

During chronic infections and cancers, CD8^+^ T cells display progressive loss of effector function accompanied with enhanced expression of inhibitory receptors and poor memory recall responses.^1,2^ These functional impairments are widely known as T cell exhaustion and are postulated as an evolutionarily conserved adaptation mechanism to persistent antigen stimulation, thereby limiting immunopathology or autoreactivity.^3,4^ Despite their altered state, exhausted T (Tex) cells still provide some protection against viral replication or tumor growth^5–7^ It is now evident that T cell exhaustion is an unique state of T cell differentiation, which is epigenetically distinct from effector T cells and memory T cells generated upon acute infection or vaccination.^8,9^ It is therefore essential to elucidate the role of epigenetic regulators governing T cell exhaustion.

Tex cells are a heterogenous population composed of functionally distinct subsets, which reflects the complex state and regulation of T cell exhaustion.^2,10–12^ Seminal studies demonstrated that Tex cells are compartmentalized into at least two major subsets: a self-renewing progenitor subset (Tex^prog^ subset), defined by high expression levels of the transcription factor T cell factor 1 (TCF1) as well as of the surface markers Ly108 and CXCR5, and a terminally exhausted TCF1^lo^ subset (Tex^term^) which is continuously replenished by Tex^prog^ cells.^13–16^ Importantly, Tex^prog^ cells display a proliferation burst upon immune checkpoint blockades (ICBs), making them an attractive and promising target for immunotherapies.^17–19^ Subsequent studies identified additional Tex cell subsets and several groups have independently identified a CX3CR1^+^ effector-like subset. This subset, designated as Tex^eff-like^ subset, retains a certain degree of cytolytic activity and thereby plays a central role in controlling chronic viral infection and tumor growth.^20–23^ Tex^eff-like^ cells display a unique transcriptional and epigenetic profile that distinguishes them from Tex^prog^ and Tex^term^ subsets and, similar to Tex^term^ cells, are derived from Tex^prog^ cells. However, although several differentiation trajectories between Tex^eff-like^ and Tex^term^ cells have been postulated,^24–26^ the exact developmental paths still remain to be clarified. Notably, ICB treatment leads to the expansion of CX3CR1^+^ cells and this correlates with the better prognosis for patients with metastatic melanoma and non-small cell lung cancer (NSCLC).^27,28^ Thus, in addition to the increase of Tex^prog^ cells, directing them into Tex^eff-like^ subset might serve as a novel therapeutic approach for chronic infection and cancer.^12,29^

In this study we elucidated the role of HDAC1, a key epigenetic regulator, in the control of CD8^+^ T cell exhaustion. By using a well-established murine model of chronic viral infection, we uncovered that HDAC1 regulates the generation and maintenance of CX3CR1^+^ Tex^eff-like^ cell subset in a T cell-intrinsic manner. Moreover, HDAC1 was essential to keep viral load below a certain level. A transcriptome analysis at single cell resolution (scRNA-seq) revealed that the deletion of HDAC1 deviates the differentiation of Tex^prog^ cells into an alternative cell subset enriched in exhaustion and cytolytic signatures. Mechanistically, HDAC1 promoted the generation of Tex^eff-like^ cell subsets, at least in part, by shaping chromatin landscapes in Tex^prog^ cells to facilitate the expression of effector-like signature genes. Together, our study demonstrates that HDAC1 functions as a key regulator for viral control during chronic infection by ensuring CX3CR1^+^ Tex^eff-like^ cell subset differentiation. Thus, targeting and modulating HDAC1-regulated pathways in CD8^+^ T cells might be an exciting therapeutic strategy toward controlling chronic viral infection.

## Results

### Mice with a T cell-specific deletion of HDAC1 display impaired viral control during chronic viral infection

To dissect the role of HDAC1 in T cell exhaustion, we induced chronic infection using the lymphocytic choriomeningitis virus (LCMV) model in T cell-specific HDAC1-deficient (*Hdac1*^f/f^, *Cd4*-Cre) and corresponding wild-type control (*Hdac1*^f/f^) mice (hereafter referred to as HDAC1-cKO and WT mice, respectively).^30^ Specifically, we infected HDAC1-cKO and WT mice with the clone 13 strain of LCMV (LCMV Cl13) (Fig. 1A), which results in chronic viral infection accompanied with a temporal weight loss driven by CD8^+^ T cells.^31–33^ Upon infection, HDAC1-cKO mice displayed a similar degree of a transient weight loss over a period of four weeks in comparison to WT mice (Fig. 1B). However, the viral titer in the serum was elevated in the absence of HDAC1 at day 30 post infection (p.i.) (Fig. 1C). This indicates an essential role of HDAC1 in T cells for controlling antiviral responses.

**Fig. 1:**
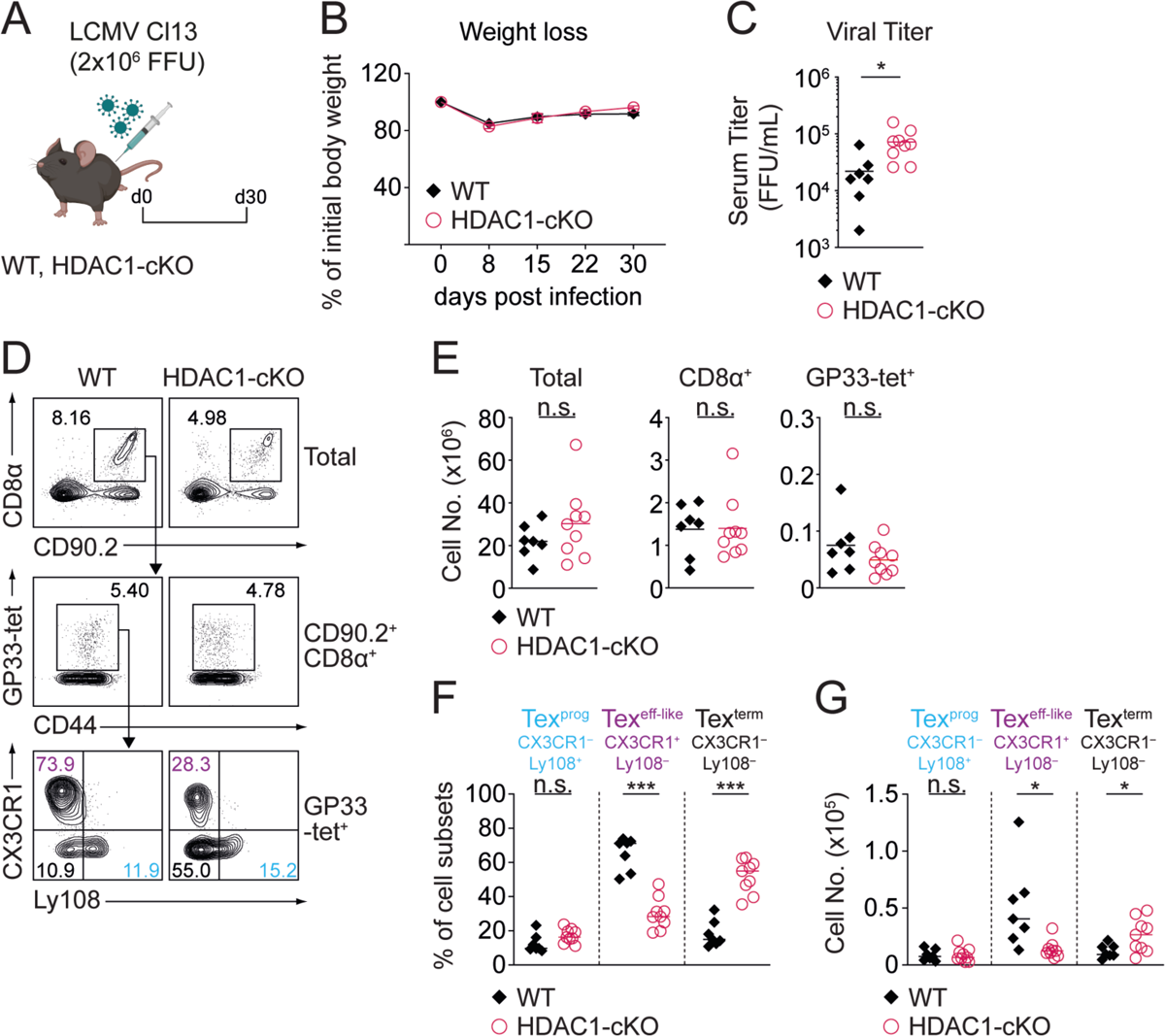
Mice with a T cell-specific deletion of HDAC1 display impaired viral control during chronic infection. (A) Schematic drawing of the experimental design. WT and HDAC1-cKO mice were infected with LCMV Cl13 and splenocytes were analyzed 30 days (d30) post infection (p.i.). (B) Diagram shows weight loss in percentage of initial body weight during the course of infection. (C) Diagram depicts viral titer in the serum of infected mice on d30 p.i. (D) Upper panel: Contour plots show percentage of WT and HDAC1-cKO CD8^+^ T cells (CD90.2^+^CD8α^+^) from total splenocytes. Middle panel: CD8^+^ T cells were further gated on CD44^+^GP33-tet^+^ viralspecific exhausted CD8^+^ T (Tex) cells. Lower panel: Virus-specific Tex cell subsets were characterized by surface expression of CX3CR1 and Ly108: Tex^prog^ (CX3CR1^-^Ly108^+^) (blue), Tex-^eff-like^ (CX3CR1^+^Ly108^-^) (purple) and Tex^term^ (CX3CR1^-^Ly108^-^) (black). (E) Diagrams show the summary of absolute numbers of total splenocytes (left), CD8α^+^ T cells (middle) and Tex cells (GP33-tet^+^) (right). (F,G) Diagrams show summary of the percentages (F) and absolute cell numbers (G) of Tex cell subsets as depicted in (D). Numbers in the plots (D) represent the percentage of cells within the indicated regions. Data are representative (D) or show the summary (B,C,E,F,G) of 7-9 mice per genotype analyzed in 2 independent experiments. (C,E,F,G) Horizontal bars indicate the mean. \**p* < 0.05, \*\**p* < 0.01, \*\*\**p* < 0.001, n.s. *p* ≥ 0.05. An unpaired two-tailed Student’s *t*-test was used to compare WT vs HDAC1-cKO cells.

Based on the expression of CX3CR1 and Ly108, three exhausted CD8^+^ T cell subsets have been defined once chronic infection has been established (> 3 weeks p.i.): Tex^prog^ cells (Ly108^+^CX3CR1^-^), Tex^eff-like^ cells (Ly108^-^CX3CR1^+^) and Tex^term^ cells (Ly108^-^CX3CR1^-^).^23^ Although there was no alteration in the numbers of splenocytes as well as total and viral glycoprotein 33-41 (GP33)-specific CD8^+^ T cells in the absence of HDAC1 (Fig. 1D and E), a detailed flow cytometry analysis of CD8^+^ T cells revealed that loss of HDAC1 led to a reduction in Tex^eff-like^ cell frequencies and numbers (Fig. 1D, F and G). This was concurrent with an expansion of the Tex^term^ subset (Fig. 1D, F and G). Moreover, given the vital role of Tex^eff-like^ cells for virus control,^21,23^ the reduction of this subset is in line with the observed increase in viremia in HDAC1-cKO mice (Fig. 1C). Together, these data indicate an essential role of HDAC1 in regulating Tex cell subset distribution and in keeping viral loads below a certain level.

### HDAC1 is essential for CX3CR1^+^ Tex cell differentiation at the onset of chronic viral infection

The emergence of Tex cells, based on transcriptional and chromatin accessibility profiling, has been observed as early as day 8 p.i..^24,25,34^ Having uncovered that HDAC1 deletion resulted in a reduction of Tex^eff-like^ cells 30 days p.i., we next aimed to determine the time point at which the differentiation of Tex cell subsets starts to be affected by HDAC1-deficiency during the course of an infection. For this we isolated peripheral blood lymphocytes from LCMV Cl13-infected HDAC1-cKO and WT mice on day 8, 15, 22 and 30 p.i. and assessed Tex cell subset distribution among GP33-tetramer^+^ (GP33-tet^+^) CD8^+^ T cells (Fig. S1A-C). Loss of HDAC1 led to a strong reduction in the frequencies of circulating CD8^+^ and GP33-tet^+^ CD8^+^ T cells (Fig. S1B). In addition, whereas WT mice displayed a strong induction of CX3CR1^+^ cells among GP33-tet^+^ CD8^+^ T cells on 8 days p.i., which is in agreement with published studies,^23,24^ the frequency of such cells was much lower in HDAC1-cKO mice on day 8, 15 and 22 (Fig. S1C).

Thus, HDAC1 deletion impaired CX3CR1^+^ cell differentiation in blood already at the onset of chronic viral infection. Ly108^+^CX3CR1^-^ cells acquire a certain degree of Tex^prog^ features as early as 8 days p.i..^23–25,35^ Moreover, fate-mapping experiments demonstrated that CX3CR1^+^ Tex cells that emerge around 8 days p.i. later give rise to a substantial proportion of a Tex^eff-like^ cell subset^22^. Therefore, we refer to these subsets on day 8 as early-Tex^prog^ cells and early-Tex^eff-like^ cells throughout the manuscript, respectively. However, given that virtually no Tex^term^ cells emerge on day 8 p.i.,^23–25,35^ we named the Ly108^-^CX3CR1^-^ subset, which characteristics are not defined (nd), as early-Tex^nd^ cells.

We next examined the composition of Tex cell subsets in the spleen of WT and HDAC1-cKO mice on day 8 p.i. (Fig. 2A). Loss of HDAC1 led to a significant reduction in CD8^+^ and GP33-tet^+^ CD8^+^ T cells (Fig. 2B and C), similar to the observation made in the blood. Additionally, the distribution of Tex cell subsets was altered in the absence of HDAC1, marked by a dramatic reduction in the frequency and number of early-Tex^eff-like^ cells (Fig. 2D-F). A comparable reduction of the early-Tex^eff-like^ cell subset was also observed for HDAC1-deficient CD8^+^ T cells specific for the viral glycoprotein 276-286 (GP276), indicating that the altered differentiation of Tex cell subsets occurs independently of TCR-specificity (Fig. S2A-C). Moreover, a severely diminished early-Tex^eff-like^ cell population was also observed in other lymphoid (i.e. inguinal LNs) as well as non-lymphoid (i.e. liver) organs in HDAC1-cKO mice (Fig. S2D and E), indicating phenotypic alterations of Tex cell subsets across all tissues. Finally, HDAC1-cKO mice also displayed a higher viral load in their serum on day 8 p.i. when compared to WT mice (Fig. 2G). This indicates that early-Tex^eff-like^ cells also play an essential role for controlling viral replication, similar to Tex^eff-like^ cells that emerge at later time points during infection.

**Fig. 2:**
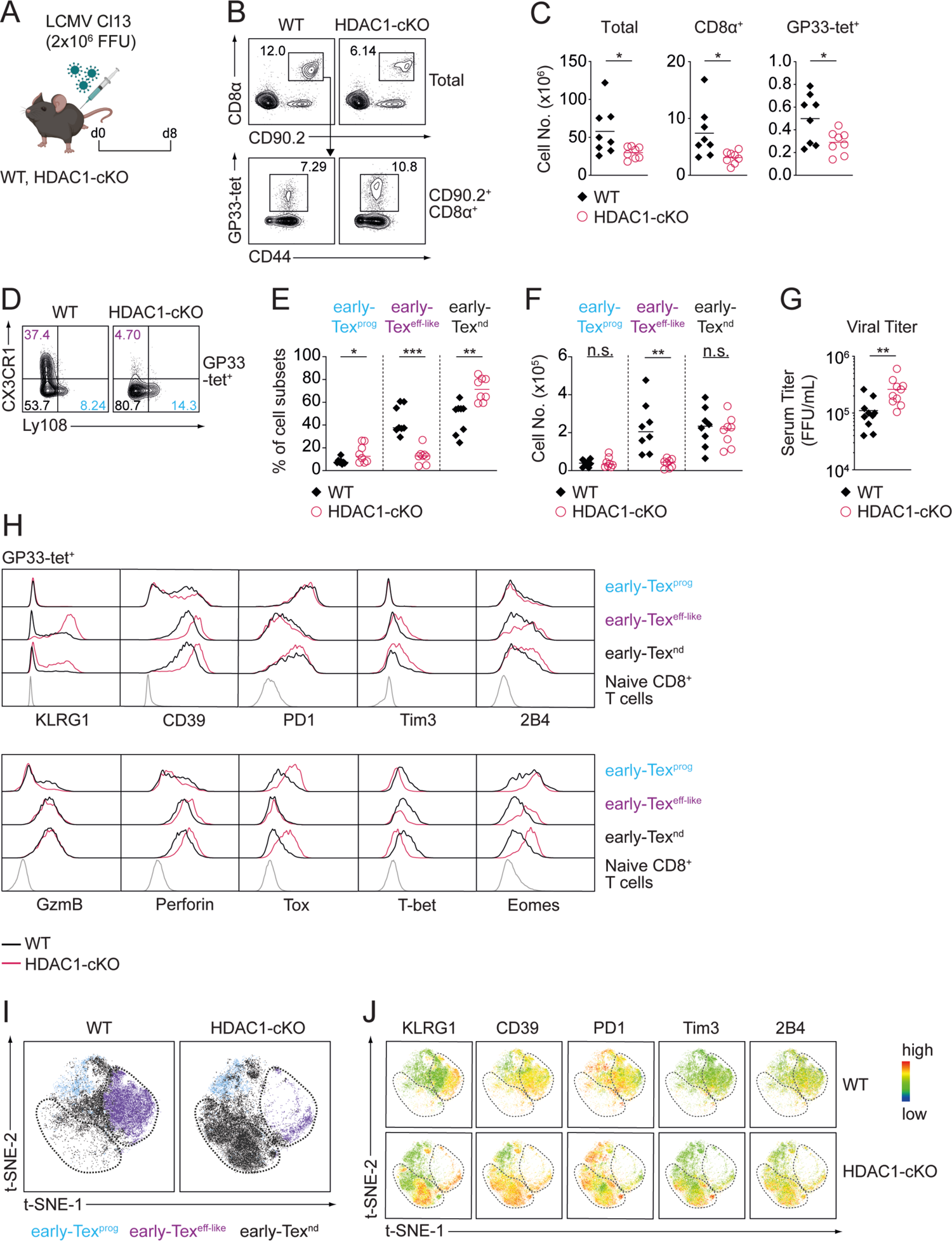
HDAC1 is essential for CX3CR1^+^ Tex cell differentiation at the onset of chronic viral infection. (A) Experimental design. WT and HDAC1-cKO mice were infected with LCMV Cl13 and splenocytes were analyzed 8 days (d8) post infection (p.i.). (B) Flow cytometry analysis showing the percentage of WT and HDAC1-cKO CD8^+^ T cells (CD90.2^+^CD8α^+^) from total splenocytes (upper panel). CD8^+^ T cells were further gated on viral-specific exhausted CD8^+^ T (Tex) cells (CD44^+^GP33-tet^+^) (lower panel). (C) Diagrams show the summary of absolute numbers of total splenocytes (left), CD8α^+^ T cells (middle) and Tex cells (GP33-tet^+^) (right). (D) Representative plots show the distribution of Tex cell subsets on d8 p.i.: early-Tex^prog^ (CX3CR1^-^Ly108^+^) (blue), early-Tex^eff-like^ (CX3CR1^+^Ly108^-^) (purple) and early-Tex^nd^ (CX3CR1^-^ Ly108^-^) (black). (E,F) Diagrams show summary of the percentages (E) and absolute numbers (F) of Tex cell subsets as depicted in (D). (G) Diagram depicts viral titer in the serum of infected mice on d8 p.i. Numbers in plots (B,D) represent the percentage of cells within the indicated regions. (H) Histograms show the expression of the surface receptors KLRG1, CD39, PD1, Tim3 and 2B4 (upper panel), and the cytolytic molecules granzyme B (GzmB) and Perforin together with the transcription factors Tox, T-bet and Eomes (lower panel) in all Tex cell subsets (pre-gated on CD8α^+^CD90.2^+^CD44^+^GP33-tet^+^ cells) on d8 p.i. (I) FlowJo generated t-SNE plots show clustering of WT and HDAC1-cKO Tex cell subsets on d8 p.i. (J) Relative expression of cell surface receptors KLRG1, CD39, PD1, Tim3 and 2B4 used as parameters for t-SNE dimensionality reduction as depicted in (I). Data are representative (B,D,H) or show the summary (C,E,F,G) of 8 mice per genotype analyzed in 2 independent experiments. Data shows summary (I,J) of concatenated samples from 4 mice per genotype. (C,E,F,G) Horizontal bars indicate the mean. \**p* < 0.05, \*\**p* < 0.01, \*\*\**p* < 0.001, n.s. *p* ≥ 0.05. An unpaired two-tailed Student’s *t*-test was used to compare WT vs HDAC1-cKO cells.

In addition to the markers Ly108 and CX3CR1, Tex cell subsets also display specific and distinct expression patterns of inhibitory receptors (IRs), effector molecules and transcription factors.^20,21,23^ Additional immunophenotyping revealed that the expression levels of IRs, such as Tim3 and 2B4 as well as of CD39, an ecto-ATPase linked with T cell exhaustion, were increased in HDAC1-cKO early-Tex^eff-like^ and early-Tex^nd^ cells (Fig. 2H, S2F and G). This was accompanied with an elevated expression of thymocyte selection-associated HMG box (Tox), a central transcription factor initiating T cell exhaustion (Fig. 2H; Fig. S2I).^34,36–39^ Moreover, loss of HDAC1 resulted in elevated Eomesodermin (Eomes) expression levels, concurrent with the downmodulation of T-Box Transcription Factor 21 (T-bet) expression (Fig. 2H; Fig. S2I). Similar expression patterns of Eomes and T-bet were shown to be linked with terminal exhaustion.^40,41^ Besides the increased expression of proteins linked to an exhaustion state, HDAC1-deficient Tex cell subsets displayed enhanced expression of Killer Cell Lectin Like Receptor G1 (KLRG1), a canonical marker for effector T cells (Fig. 2H; Fig. S2F).^10,42,43^ In addition, while there was no change in granzyme B (GzmB) expression, perforin expression was higher in HDAC1-deficient early-Tex^eff-like^ and early-Tex^nd^ cells (Fig. 2H; Fig. S2H), indicating that certain effector features are enhanced upon loss of HDAC1. However, the expression of exhaustion-related factors (Fig. 2H; Fig. S2F-I) and the reduced number of Tex cells (Fig. 2C) during infection might not allow proper viral clearance in these mice, resulting in increased viremia (Fig. 2G). To gain further insight into the impact of HDAC1 deletion on the heterogeneity and subset composition of Tex cells, we applied t-SNE dimensionality reduction to our flow cytometry data. While a population corresponding to early-Tex^eff-like^ cells was easily detectable in WT CD8^+^ T cells (Fig. 2I, encircled purple population), deletion of HDAC1 led to an almost complete loss of such a subset (Fig. 2I). Instead, a unique population (encircled black population) co-expressing IRs and KLRG1 at high levels emerged in HDAC1-cKO mice (Fig. 2I and J). Taken together, these results highlight an essential role of HDAC1 in promoting the generation of early-Tex^eff-like^ cells.

### HDAC1 controls early-Tex^eff-like^ subset differentiation in a CD8^+^ T cell-intrinsic manner

The formation of Tex^eff-like^ cells requires help from CD4^+^ T cells that secrete interleukin-21 (IL-21).^22,23^ Since *Hdac1* is deleted both in CD4^+^ and CD8^+^ T cells in HDAC1-cKO mice (owing to the expression pattern of Cre recombinase in the *Cd4*-Cre deleter line),^44^ alterations in the subset distribution of Tex cells in these mice might be due to a dysfunction of either CD4^+^ or CD8^+^ T cells, or both. To distinguish among those possibilities, we performed adoptive transfer experiments with CD8^+^ T cells that express the transgenic P14 TCR, recognizing viral GP33 peptide presented by H2D^b^.^45^ We crossed WT and HDAC1-cKO mice with P14 TCR transgenic mice and additionally introduced a *Rosa26*-STOP-EYFP reporter allele^46^ (hereafter referred to as P14-WT and P14-HDAC1-cKO mice, respectively). This allows the tracking of YFP^+^ P14-HDAC1-cKO T cells due to Cre-mediated deletion of the STOP cassette (Fig. 3A). Naïve P14-WT (CD90.2^+^) and P14-HDAC1-cKO (CD90.2^+^YFP^+^) cells were mixed at a 1:1 ratio and adoptively co-transferred into wild-type CD90.1^+^ recipient mice. On the following day recipient mice were infected with LCMV Cl13 and the differentiation of Tex cell subsets was analyzed 8 days p.i. (Fig. 3B). Consistent with the reduced number of virus-specific CD8^+^ T cells observed in HDAC1-cKO mice (Fig. 2C), the proportion of P14-HDAC1-cKO cells was reduced in a competitive setting (Fig. 3C and D). Moreover, P14-HDAC1-cKO cells “phenocopied” the distribution of Tex cell subsets that has been observed in HDAC1-cKO mice (Fig. 2D-F), showing reduced frequencies of early-Tex^eff-like^ cells (Fig. 3E and F). Of note, we also generated mixed bone barrow (BM) chimeric mice, where WT (CD45.2^+^) or HDAC1-cKO (CD45.2^+^) BM cells were mixed at a 1:1 ratio with congenically distinguishable wild-type (CD45.1^+^) BM cells (Fig. S3A). Following reconstitution of irradiated recipients and subsequent infection, we observed reduced frequencies of the early-Tex^eff-like^ cell subset within the reconstituted CD45.2^+^ HDAC1-deficient but not CD45.2^+^ WT compartment (Fig. S3B-C). Together, these results clearly indicate that HDAC1 has a key and CD8^+^ T cell-intrinsic role in the generation of early-Tex^eff-like^ subsets.

**Fig. 3:**
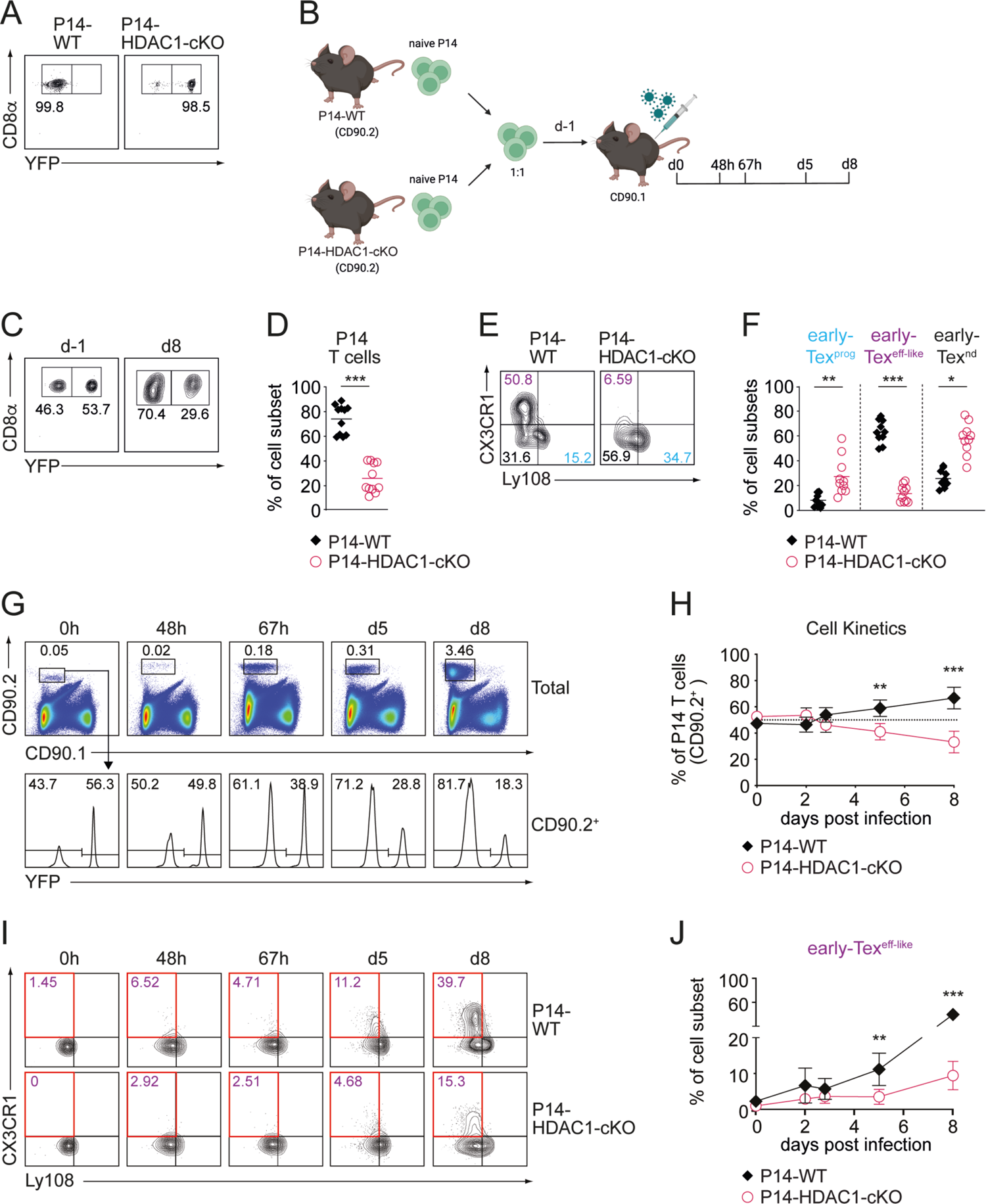
HDAC1 controls early-Tex^eff-like^ subset differentiation in a CD8^+^ T cell-intrinsic manner. (A) Contour plots show YFP and CD8α expression in freshly isolated naïve P14-WT and P14-HDAC1-cKO cells. (B) Scheme of the experimental design. Naïve P14-WT and P14-HDAC1-cKO CD8^+^ T cells (CD90.2^+^) were isolated and transferred into congenic CD90.1^+^ recipient mice one day prior (d-1) to LCMV Cl13 infection. The differentiation kinetics and the composition of Tex cell subsets were determined at various time points after infection (0h, 48h, 67h, d5 and d8). (C) Contour plots show the percentages of YFP^-^ P14-WT and YFP^+^ P14-HDAC1-cKO cells on the day of transfer (d-1) and on d8 post infection (p.i.). (D) Diagram shows the summary of the percentages of transferred P14-WT and P14-HDAC1-cKO cells on d8 p.i. as depicted in (C). (E) Contour plots show the distribution of Tex cell subsets on d8 p.i.: Tex^prog^ (CX3CR1^-^Ly108^+^) (blue), early-Tex^eff-like^ (CX3CR1^+^Ly108^-^) (purple) and early-Tex^nd^ (CX3CR1^-^Ly108^-^) (black) within transferred P14 cells. (F) Diagrams show summary of the percentages of Tex cell subsets as depicted in (E). (G) Density plots show percentages of transferred P14 CD8^+^ T cells (CD90.2^+^) in the spleen of infected mice at various time points of infection (0h, 48h, 67h, d5 and d8) (upper panel). P14-WT and P14-HDAC1-cKO cells were distinguished by the expression of YFP (lower panel). (H) Summary diagram shows percentages of transferred YFP^-^ P14-WT and YFP^+^ P14-HDAC1-cKO CD8^+^ T cells at various time points p.i. as depicted in (G). (I) Contour plots show the distribution of P14-WT and P14-HDAC1-cKO Tex cell subsets at various time points of infection. (J) Summary diagram shows percentage of early-Tex^eff-like^ subsets as depicted in (I, red square). Numbers in the plots (A,C,E,G,I) show the percentage of cells within the indicated regions. Data shown are representative (A,C,E,G,I) or show the summary (D,F,H,J) of 10 mice per genotype analyzed in 3 independent experiments (D,F) or of 3-8 mice per time point per genotype from at least 3 independent experiments (H,J). (D,F) Horizontal bars indicate the mean. (H,J) Data are shown as mean ± SD. (D,F,H,J) An unpaired (D,F) or paired (H,J) two-tailed Student’s *t*-test was used to compare P14-WT vs P14-HDAC1-cKO cells. \**p* < 0.05, \*\**p* < 0.01, \*\*\**p* < 0.001, n.s. *p* ≥ 0.05.

### HDAC1-mediated control of early-Tex^eff-like^ subset differentiation is not due to a defect in early CD8^+^ T cell activation, proliferation or survival

Upon the onset of chronic infection, viral-specific CD8^+^ T cells are activated, clonally expand and differentiate into Tex cell subsets.^24,35,42^ In order to define at which timepoint upon infection Tex cell subset composition is altered in the absence of HDAC1, we examined the kinetics of the appearance of CX3CR1^+^ cells at day 3, 5 and 8 p.i., and also determined the activation and expansion of early-Tex cells. We employed the aforementioned adoptive P14 T cell co-transfer model (Fig. 3B) and observed no change in the ratio of transferred WT to HDAC1-cKO P14 T cells up to 67 hours p.i. (Fig. 3G and H). In addition, there was no difference in cell size (as revealed by forward scatter values) as well as in the expression levels of the early activation marker CD69 between the two groups at the onset of activation (Fig. S4A and B). Furthermore, both WT and HDAC1-deficient cells divided at a similar rate based on the “dilution” of division-tracking dye CellTrace Violet (CTV) (Fig. S4A and B). These data indicate intact TCR-signaling in the absence of HDAC1 at early timepoints. In contrast, from day 5 p.i. on HDAC1-deficient P14 T cells displayed a relative reduction compared to the WT cells (Fig. 3G and H). In addition, while the appearance of CX3CR1^+^ cells first became evident at day 5 p.i. in the WT compartment, the proportion of these cells was severely reduced in the absence of HDAC1 (Fig. 3I and J). However, the deletion of HDAC1 had no impact on the degree of ongoing proliferation as well as the proportion of apoptotic cells (assessed by intracellular Ki67 and active caspase-3 staining, respectively) over the assessed time period of 8 days p.i. (Fig. S4C and D). Together, these data indicate that HDAC1-mediated regulation of early-Tex^eff-like^ cell differentiation is independent of the early activation, proliferation or survival of these cells. Rather, HDAC1 might represent a checkpoint that instructs cell fate specification and differentiation towards early-Tex^eff-like^ population as early as 5 days p.i..

### HDAC1 is required for the maintenance of the Tex^eff-like^ cell pool

Our data clearly demonstrate a key role of HDAC1 in the generation of early-Tex^eff-like^ cells. Since previous studies have shown that Tex^eff-like^ cell pool is maintained throughout chronic viral infection,^23,24^ we next sought to examine whether HDAC1 is also essential for the maintenance of Tex^eff-like^ cells. In order to address this question, we crossed *Hdac1*^f/f^ mice onto a *Rosa26*-CreERT2 background^47^ to inducibly delete *Hdac1* by tamoxifen administration during the course of chronic viral infection. We also introduced a *Rosa26*-STOP-EYFP reporter allele to monitor Cre activity and thus cells that have deleted *Hdac1*, leading to the generation of *Hdac1*^f/f^, *Rosa26*-CreERT2/STOP-YFP and *Hdac1*^+/+^, *Rosa26*-CreERT2/STOP-YFP mice (hereafter referred to as WT^CreERT-YFP^ and HDAC1-cKO^CreERT-YFP^ mice, respectively). In the absence of tamoxifen, YFP expression was not induced and the distribution of circulating Tex cell subsets was normal 8 days post LCMV Cl13 infection, indicating that the genetic system is tightly controlled as expected (Fig. S4E and F). Subsequently, mice were treated with tamoxifen on day 9 p.i. and the distribution of Tex cell subsets in the spleen was analyzed 6 days later (i.e. 15 days p.i.) (Fig. 4A). After tamoxifen injection, we observed an induction of YFP expression in approx. 30% of CD8^+^ T cells. However, this didn’t correlate with the loss of HDAC1 expression, which was detected in approx. 40% of CD8^+^ T cells (Fig. S4G). Therefore, HDAC1-deficient cells in HDAC1-cKO^CreERT-YFP^ mice were identified by “gating” on the HDAC1-negative (HDAC1^-^) population based on intracellular anti-HDAC1 staining (Fig. 4B and C). Strikingly, the analysis of the distribution of Tex cell subsets revealed a reduction in the proportion of the CX3CR1^+^ cells within the HDAC1^-^ population in HDAC1-cKO^CreERT-YFP^ mice compared to WT ^CreERT-YFP^ mice as well as to the HDAC1^+^ population of HDAC1-cKO^CreERT-YFP^ mice (Fig. 4D and E). Taken together, our results demonstrate a CD8^+^ T cell-intrinsic key role for HDAC1 not only for the generation but also for the maintenance of the Tex^eff-like^ cell population during chronic viral infection.

**Fig. 4:**
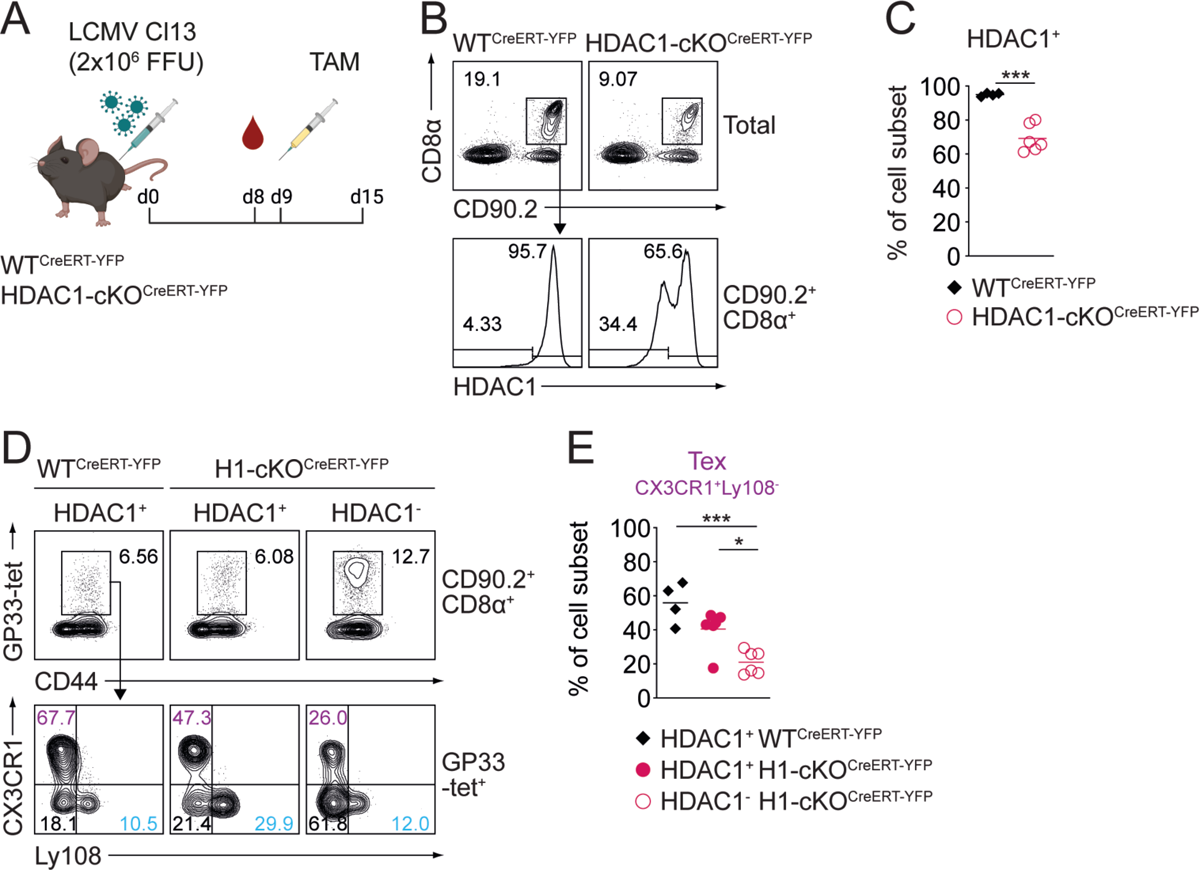
HDAC1 is required for the maintenance of the Tex^eff-like^ cell pool. (A) Experimental design. WT^CreERT-YFP^ and HDAC1-cKO^CreERT-YFP^ mice were infected with LCMV Cl13. On day 8 (d8) post infection (p.i.), the composition of Tex cell subsets was investigated in the blood of infected mice. One day later (d9), mice were injected with 2 mg of Tamoxifen (TAM) to induce the deletion of HDAC1. Splenocytes were analyzed on d15 p.i. (B) Contour plots show the percentage of WT^CreERT-YFP^ and HDAC1-cKO^CreERT-YFP^ CD8^+^ T cells (CD90.2^+^CD8α^+^) of total lymphocytes (upper panel). CD8^+^ T cells were further characterized as HDAC1-sufficient (HDAC1^+^) or HDAC1-deficient (HDAC1^-^) cells (lower panel). (C) Diagrams shows summary of percentage of HDAC1 expressing (HDAC1^+^) CD8^+^ T cells as depicted in (B). (D) Contour plots show GP33-tet^+^ Tex cells (upper panel) and the distribution of Tex cell subsets: CX3CR1^-^ Ly108^+^ Tex (blue), CX3CR1^+^Ly108^-^ Tex (purple) and CX3CR1^-^Ly108^-^ Tex (black) (lower panel) within HDAC1-sufficient (HDAC1^+^) and HDAC1-deficient (HDAC1^-^) CD8^+^ T cells. (E) Summary diagram shows the percentage of CX3XR1^+^Ly108^-^ Tex cell subset as depicted in (D). Numbers in plots (B,D) show the percentage of cells within the indicated regions. Data are representative (B,D) or show the summary (C,E) of 4-6 mice per genotype analyzed in 2 independent experiments. (C,E) Horizontal bars indicate the mean. \**p* < 0.05, \*\**p* < 0.01, \*\*\**p* < 0.001, n.s. *p* ≥ 0.05. (C,E) An unpaired two-tailed Student’s *t*-test was used to compare WT^Cre-ERT-YFP^ vs HDAC1-cKO^CreERT-YFP^ cells. (E) One-way ANOVA followed by Tukeýs multiple-comparison test was used for the comparison of more than 2 groups.

### scRNA-seq reveals transcriptionally distinct cell clusters between LCMV-specific WT and HDAC1-deficient CD8^+^ T cells

In order to gain further insight into the impact of HDAC1-deletion on Tex cell subset diversity at early-stages during chronic infection, we performed single-cell RNA sequencing (scRNA-seq) of naïve and LCMV-specific (GP33-tet^+^) CD8*^+^* T cells isolated from WT and HDAC1-cKO mice, encompassing uninfected and day 8 infected conditions, respectively. Uniform manifold approximation and projection (UMAP) followed by Seurat-based clustering led to the identification of 8 clusters (Fig. 5A and B; Table 1), which were subsequently annotated based on their unique marker genes (Fig. 5C) as well as their similarity to published signature genes of CD8^+^ T cell subsets upon acute or chronic LCMV infection (Fig. S5A-C).^24^ We found two canonical clusters corresponding to naïve (cluster C1; T^Naïve^) and early-Tex^prog^ cells (C2; Tex^prog^). Furthermore, two other relatively small clusters were identified that highly expressed genes related to either cell proliferation (e.g. *Hist1h1b, Mki67*) (C3; Tex^prol^) or terminal exhaustion (e.g. *Bcl2a1d, Lag3*) (C4; Tex^exh^) (Fig. 5A-C; Fig. S5A-C). The majority of cells (approx. 77% of WT and 67% of HDAC1-cKO cells) were grouped into four other clusters (C5-C8) (Fig. S5B. Cluster C5 (Tex^early^) was enriched for a gene signature of Tex^eeff^ cells (early effector exhausted cells where Tex program is initiated^24^) (Fig. S5A and C). Cluster C6 (Tex^int^) expressed the *Cx3cr1* gene and was enriched for a gene signature of Tex^int^ cells (intermediate exhausted cells harboring potential to become both Tex^term^ and Tex^eff-like^ cells^24^) (Fig. 5C; Fig. S5A and C). Cluster C7 (Tex*^Cx3cr1^)* was expressing the *Cx3cr1* gene at a higher level than the C6 cluster and was enriched for a gene signature of Tex^KLR^ cells (KLR family member protein-expressing Tex cells, a major subpopulation of Tex^eff-like^ cells^24^) (Fig. 5C; Fig. S5A and C). Lastly, cluster C8 (Tex^cyt^) was highly expressing genes encoding cytolytic proteins but also displaying the highest enrichment score of a Tex^term^ gene signature^24^) (Fig. 5C; Fig. S5A and C). Notably, while the C1-C4 clusters (T^Naïve^, Tex^prog^, Tex^prol^, and Tex^exh^) were almost equally composed of WT and HDAC1-cKO cells, cluster C6 (Tex^int^) and cluster C7 (Tex*^Cx3cr1^*) contained mostly WT cells (Fig. 5A and B; Fig. S5B). In contrast to the C6 and C7 clusters, the vast majority of the C5 (Tex^early^) and C8 (Tex^cyt^) clusters consisted of HDAC1-deficient cells (Fig. 5A and B; Fig. S5B). A gene ontology (GO) analysis of clusters C5-C8 revealed that several biological processes (e.g. ribonucleoprotein complex biogenesis) were uniquely overrepresented in HDAC1-cKO-dominant C5 cluster (Tex^early^) (Fig. 5D; Fig. S5D). In addition, while WT-dominant clusters C6 and C7 shared almost identical enriched terms for GO biological processes, some of these processes (e.g. “leukocyte migration”) were not enriched in HDAC1-cKO-dominant C5 and C8 (Fig. 5D; Fig. S5D). Instead, cluster C8 shared certain processes with C5 (e.g. “oxidation of organic compounds”) (Fig. 5D; Fig. S5D). This analysis suggests that the deletion of HDAC1 leads to the emergence of transcriptionally distinct subsets within the early-non-Tex^prog^ population (i.e. Tex cells other than early-Tex^prog^ cells), in contrast to its minor impact on the generation of early-Tex^prog^ cells. In line with our immunophenotyping of early-Tex cell subsets (Fig. 2D-F and H-J; Fig. S2F-I), a comparison of gene expression between WT and HDAC1-cKO early-non-Tex^prog^ cells showed that the deletion of HDAC1 led to the increased expression of genes associated with cytolytic function (e.g. *Gzma*, *Gzmk*, *Prf1*, *Klre1*) as well as exhaustion (e.g. *Havcr2, Tox, Tnfrsf9, Eomes*), concurrent with the downmodulation of *Cx3cr1* gene expression (Fig. 5E). Thus, the deletion of HDAC1 resulted in specific transcriptional alterations and the enlargement of two Tex clusters with early exhaustion (C5) and exhaustion/cytolysis (C8) signatures.

**Fig. 5:**
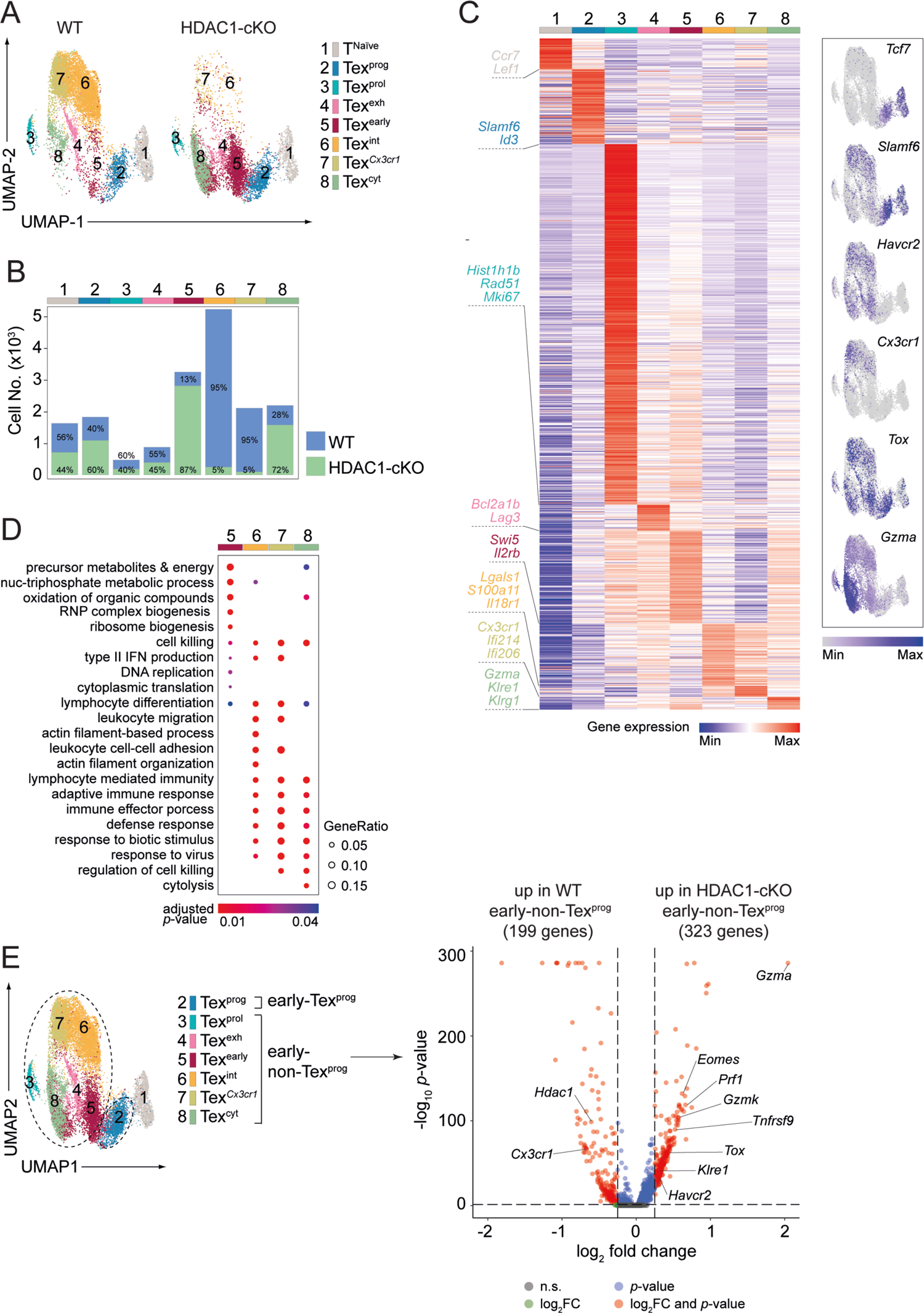
scRNA-seq reveals transcriptionally distinct cell clusters between LCMV-specific WT and HDAC1-deficient CD8^+^ T cells. (A) UMAP plots display clusters identified from a pool of naïve and virus-specific (GP33-tet^+^) WT and HDAC1-cKO T cells from uninfected and from day 8 LCMV-Cl13-infected mice, respectively. (C1) T^naive^; (C2) Tex^prog^: progenitor; (C3) Tex^prol^: proliferating; (C4) Tex^exh^: exhausted; (C5) Tex^early^: early; (C6) Tex^int^: intermediate; (C7) Tex*^Cx3cr1^*; (C8) Tex^cyt^: cytotoxic. (B) Stacked bar graph shows the number of WT or HDAC1-cKO CD8^+^ T cells within each cluster as defined in (A). Numbers in the bars indicate the frequency of WT or HDAC1-cKO CD8^+^ T cells within a particular cluster. (C) Left: heatmap showing the average expression of all genes unique for each cluster as defined by FindAllMarkers() from the Seurat package using default parameters. Selected genes representative for each cluster are shown. Right: feature plots of selected subset-specific marker genes indicating the expression level in each cell. Contrast was improved by using minimum and/or maximum cut-offs and positive cells were plotted on top. (D) Bubble plot showing significantly enriched (adjusted *p*-value < 0.05) GO terms (biological processes) in the WT (C6 and C7) and HDAC1-cKO (C5 and C8) dominant clusters obtained from clusterProfiler. Top 200 markers of each cluster were used for enrichment analysis. Top 5 enriched pathways from each cluster were collapsed to parent terms using Revigo and then visualized. Bubble size corresponds to the ratio of marker genes overlapping with the given pathway (Gene Ratio), bubble color corresponds to the adjusted *p*-value. (E) Left: UMAP shows the defined early-Tex^prog^ cell cluster (C2) and the early-non-Tex^prog^ cell cluster (C3-8). Right: volcano plot showing differentially expressed genes (DEGs) between WT and HDAC1-cKO early-non-Tex^prog^ cells. Horizontal dashed line at *p-*value of 0.05, vertical dashed lines at log_2_FC of −0.25 and 0.25, FC: fold change, n.s.: not significant. Numbers on plots represent the cluster number as assigned by scRNA-seq analysis (A,E) or the frequencies within the indicated regions (B).

**Table 1:** Cell clusters and frequencies of LCMV-specific WT and HDAC1-deficient CD8^+^ T cells as defined by scRNA-seq.

| Cluster Number | Cluster Name | Genotype | Cell Number | Frequency of Genotype within Cluster | Frequency of Cluster within Genotype |
| --- | --- | --- | --- | --- | --- |
| 1 | T<sup>Naive</sup> | WT | 855 | 56% | 8.8% |
| 2 | Tex<sup>prog</sup> | WT | 688 | 40% | 7.1% |
| 3 | Tex<sup>prol</sup> | WT | 267 | 60% | 2.7% |
| 4 | Tex<sup>exh</sup> | WT | 450 | 55% | 4.6% |
| 5 | Tex<sup>early</sup> | WT | 404 | 13% | 4.1% |
| 6 | Tex<sup>int</sup> | WT | 4,640 | 95% | 47.6% |
| 7 | Tex<sup>Cx3cr1</sup> | WT | 1,873 | 95% | 19.2% |
| 8 | Tex<sup>cyt</sup> | WT | 572 | 28% | 5.9% |
| 1 | T<sup>Naive</sup> | HDAC1-cKO | 664 | 44% | 10.0% |
| 2 | Tex<sup>prog</sup> | HDAC1-cKO | 1,015 | 60% | 15.3% |
| 3 | Tex<sup>prol</sup> | HDAC1-cKO | 176 | 40% | 2.6% |
| 4 | Tex<sup>exh</sup> | HDAC1-cKO | 367 | 45% | 5.5% |
| 5 | Tex<sup>early</sup> | HDAC1-cKO | 2,624 | 87% | 39.5% |
| 6 | Tex<sup>int</sup> | HDAC1-cKO | 233 | 5% | 3.5% |
| 7 | Tex<sup>Cx3cr1</sup> | HDAC1-cKO | 93 | 5% | 1.4% |
| 8 | Tex<sup>cyt</sup> | HDAC1-cKO | 1,472 | 72% | 22.2% |

### HDAC1 alters the chromatin landscape at signature gene loci associated with early-Tex-eff-like cells

Transcriptional changes during Tex subset differentiation are associated with dynamic shifts in chromatin accessibility.^20,48^ Given that our data shows no alteration in the activation, proliferation and survival of early-Tex cells (Fig. S4A-D) as well as the intact generation of early-Tex^prog^ cells by loss of HDAC1 (Fig. 5A and B), it is conceivable that HDAC1 controls the transition process of early-Tex^prog^ cells into early-Tex^eff-like^ cells, at least in part, by mediating changes in the chromatin landscape. To examine this hypothesis, we conducted bulk assay for transposase accessible chromatin sequencing (ATAC-seq) of P14-WT and P14-HDAC1-cKO early-Tex^prog^ (Ly108^+^Tim3^-^) as well as early-non-Tex^prog^ (Ly108^-^Tim3^+^) cells 8 days post LCMV Cl13 infection. A comparison between WT and HDAC1-cKO cells revealed 319 and 2,179 differentially accessible regions (DARs) in early-Tex^prog^ and early-non-Tex^prog^ cells, respectively (Fig. 6A and B; Table 2). The vast majority of the DARs were located at introns, intergenic regions or promoters in both populations, and loss of HDAC1 generally led to increased numbers of “open” chromatin regions in both populations, in line with its function in promoting a “closed” chromatin structure (Fig. 6A and B). We next explored whether there is a correlation between changes in chromatin accessibility and gene expression by integrating the data obtained from bulk ATAC-seq and scRNA-seq. For this, we defined the gene sets harboring DARs either in WT or HDAC1-cKO cells and determined the overall relative expression levels (i.e. the module scores) of these gene sets at single-cell level (Fig. 6C; Fig. S6A). Moreover, we calculated the average module scores within the individual clusters (i.e. C2-C8) and thereby examined whether DARs-associated genes are preferentially expressed in certain cluster(s) (Fig. 6D; Fig. S6B). Notably, this analysis revealed that genes more accessible in WT early-Tex^prog^ cells are highly expressed in WT-dominant cluster C6 (Tex*^int^*) and C7 (Tex*^Cx3cr1^*), indicating a role for HDAC1 in priming the expression of early-Tex^eff-like^ cell-associated genes by “opening” their loci in early-Tex^prog^ cells (Fig. 6C and D). In contrast, for genes associated with DARs more accessible in HDAC1-deficient early-Tex^prog^ cells, there was no enrichment to a particular cluster (Fig. 6C and D). Overrepresentation analysis of GO terms showed that genes associated with DARs more accessible in WT early-Tex^prog^ cells are also enriched in GO terms such as “leukocyte cell-cell adhesion” and “lymphocyte differentiation” (Fig. S6C), similar to the single-cell transcriptome of WT-dominant clusters C6 and C7 (Fig. 5D). Moreover, gene set enrichment analysis (GSEA) of DAR-associated genes revealed the overrepresentation of an effector T cell signature (i.e. a signature of total CD8^+^ T cells 8 days post acute LCMV infection^49^) in WT cells in comparison to the mutant cells (Fig. 6E). These results further support a role for HDAC1 in priming an early-Tex^eff-like^ program. Of note, genes more accessible in WT or HDAC1-cKO early-non-Tex^prog^ cells were highly expressed in WT- or HDAC1-cKO-dominant clusters (i.e. C6/C7 or C5/C8), respectively, indicating a correlation of accessibility and gene expression in early-non-Tex^prog^ cells (Fig. S6A and B). Finally, an enrichment analysis of transcription factor (TF)-binding motifs revealed shared as well as unique TF-binding motifs enriched in DARs of WT and HDAC1-cKO cells, both in early-Tex^prog^ cells as well as in the early-non-Tex^prog^ cell population (Fig. 6F; Fig. S6D). This suggests that the same TFs might be recruited to different gene loci and that a different set of TFs might be active, dependent on the presence or absence of HDAC1. Together, the integration of our ATAC-seq analysis with our scRNA-seq data indicates that HDAC1 is essential for opening chromatin at effector-like signature gene loci in early-Tex^prog^ cells. Thus, HDAC1 primes the expression of genes associated with early-Tex^eff-like^ cells, thereby promoting early-Tex^eff-like^ cell differentiation.

**Fig. 6:**
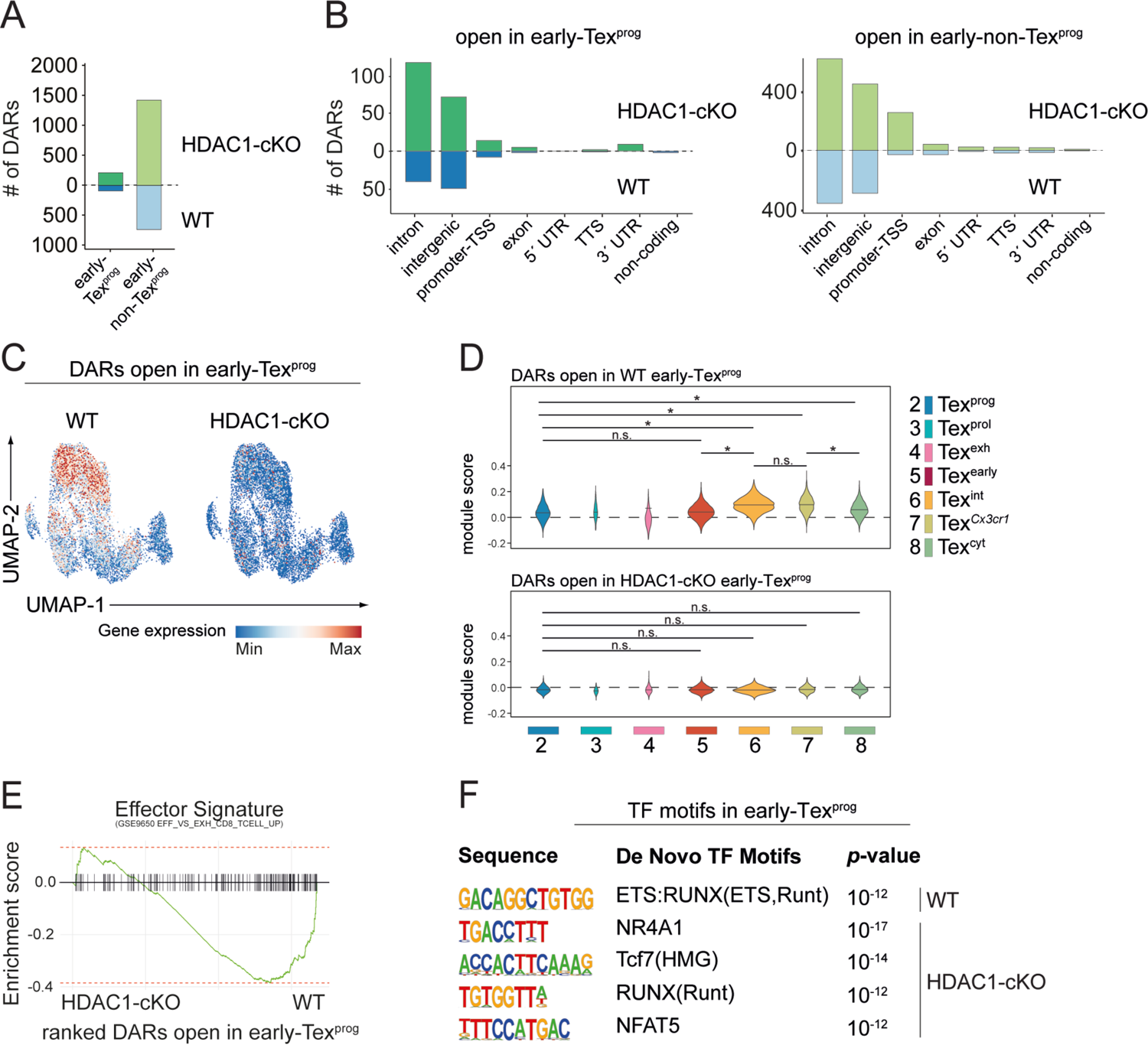
HDAC1 alters the chromatin landscape at signature gene loci associated with early-Tex^eff-like^ cells. (A) Diagram indicates the number of differentially accessible regions (DARs) (adjusted *p*-value < 0.05) between WT and HDAC1-cKO early-Tex^prog^ and early-non-Tex^prog^. (B) Number of significantly DARs in early-Tex^prog^ (left) and early-non-Tex^prog^ (right) split by their detailed genomic annotation obtained by HOMER. TSS: transcription start site (−1 kb to +100 bp around TSS), TTS: transcription termination site (−100 bp to +1 kb around TTS) (C) UMAP plot depicting the expression of genes (as revealed by scRNA-seq) associated with open DARs in either WT (left) or HDAC1-cKO (right) early-Tex^prog^ from ATAC-seq. (D) Violin plots depict the average expression of genes associated with open DARs in WT early-Tex^prog^ cells (upper panel) or HDAC1-cKO early-Tex^prog^ cells (lower panel), calculated as module scores for the clusters C2-C8 as defined by scRNA-seq. ANOVA followed by Tukey’s test was performed, * adjusted *p*-value < 0.05, n.s. not significant. (E) Gene set enrichment analysis (GSEA) shows Effector Signature (from ImmuneSigDB) of ranked genes associated with DARs open in either WT or HDAC1-cKO early-Tex^prog^ cells. The barcodes indicate the location of the genes associated with DARs in the ranked list which intersect with the respective signature. (F) Transcription factor (TF) motifs within DARs open in either WT or HDCA1-cKO early-Tex^prog^ cells as determined by de novo motif analysis using HOMER (*p*-values < 10^-11^ were considered true positive).

**Table 2:** Number of DARs in WT and HDAC1-deficient early-Tex^prog^ and early-non-Tex^prog^ as defined by ATAC-seq.

| Group | Genotype | Annotation | Number of DARs |
| --- | --- | --- | --- |
| Tex<sup>prog</sup> | WT | intron | 40 |
| Tex<sup>prog</sup> | WT | intergenic | 49 |
| Tex<sup>prog</sup> | WT | promoter-TSS | 8 |
| Tex<sup>prog</sup> | WT | exon | 2 |
| Tex<sup>prog</sup> | WT | 5' UTR | 0 |
| Tex<sup>prog</sup> | WT | TTS | 1 |
| Tex<sup>prog</sup> | WT | 3' UTR | 0 |
| Tex<sup>prog</sup> | WT | non-coding | 2 |
| Tex<sup>prog</sup> | WT | total | 102 |
| Tex<sup>prog</sup> | HDAC1-cKO | intron | 116 |
| Tex<sup>prog</sup> | HDAC1-cKO | intergenic | 71 |
| Tex<sup>prog</sup> | HDAC1-cKO | promoter-TSS | 14 |
| Tex<sup>prog</sup> | HDAC1-cKO | exon | 5 |
| Tex<sup>prog</sup> | HDAC1-cKO | 5' UTR | 0 |
| Tex<sup>prog</sup> | HDAC1-cKO | TTS | 2 |
| Tex<sup>prog</sup> | HDAC1-cKO | 3' UTR | 9 |
| Tex<sup>prog</sup> | HDAC1-cKO | non-coding | 0 |
| Tex<sup>prog</sup> | HDAC1-cKO | total | 217 |
| non-*Tex*<sup>prog</sup> | WT | intron | 358 |
| non-*Tex*<sup>prog</sup> | WT | intergenic | 290 |
| non-*Tex*<sup>prog</sup> | WT | promoter-TSS | 29 |
| non-*Tex*<sup>prog</sup> | WT | exon | 29 |
| non-*Tex*<sup>prog</sup> | WT | 5' UTR | 7 |
| non-*Tex*<sup>prog</sup> | WT | TTS | 18 |
| non-*Tex*<sup>prog</sup> | WT | 3' UTR | 15 |
| non-*Tex*<sup>prog</sup> | WT | non-coding | 3 |
| non-*Tex*<sup>prog</sup> | WT | total | 749 |
| non-*Tex*<sup>prog</sup> | HDAC1-cKO | intron | 618 |
| non-*Tex*<sup>prog</sup> | HDAC1-cKO | intergenic | 447 |
| non-*Tex*<sup>prog</sup> | HDAC1-cKO | promoter-TSS | 255 |
| non-*Tex*<sup>prog</sup> | HDAC1-cKO | exon | 41 |
| non-*Tex*<sup>prog</sup> | HDAC1-cKO | 5' UTR | 23 |
| non-*Tex*<sup>prog</sup> | HDAC1-cKO | TTS | 21 |
| non-*Tex*<sup>prog</sup> | HDAC1-cKO | 3' UTR | 18 |
| non-*Tex*<sup>prog</sup> | HDAC1-cKO | non-coding | 7 |
| non-*Tex*<sup>prog</sup> | HDAC1-cKO | total | 1430 |

## Discussion

Understanding transcriptional and epigenetic mechanisms underlying Tex cell subset differentiation will provide profound insight for the development of innovative immunotherapies against chronic infection and tumors.^3,8,9,12^ Here, we discovered a crucial role of HDAC1 for the generation and maintenance of effector-like subsets during chronic viral infection. T cell-specific HDAC1-deficient mice displayed a substantial loss of CX3CR1^+^ Tex^eff-like^ cells during the course of LCMV Cl13 infection, correlating with elevated viremia. Single-cell transcriptome analysis revealed that the generation of early-Tex^prog^ cells was largely intact in the absence of HDAC1. However, HDAC1-deficient early-Tex^prog^ cells deviated into alternative cell subsets that emerged in WT mice only at low frequencies, including a subset enriched with both exhaustion and cytolytic features. Mechanistically, HDAC1 promoted the differentiation of early-Tex^prog^ cells into early-Tex^eff-like^ cell subsets, in part by changing the chromatin landscape in early-Tex^prog^ cells that primes the expression of Tex^eff-like^ cell subset associated genes. Thus, HDAC1 is an integral part of the differentiation program endowing Tex cells with effector-like characteristics.

Current models of Tex differentiation suggest that both Tex^eff-like^ and Tex^term^ cell subsets are generated from Tex^prog^ cells.^20,21,23^ However, little is known about the molecular mechanisms directing Tex^eff-like^ or Tex^term^ cell subsets.^12,29^ Based on our data showing a reduction of Tex^eff-like^ cells in the absence of HDAC1, HDAC1 might promote early-Tex^prog^ towards early-Tex^eff-like^ differentiation or it might suppress alternative early-Tex^prog^ fates. The deletion of HDAC1 increased the accessibility of certain gene loci in early-Tex^prog^ cells, as one might have expected from the well-known function of HDACs to promote a closed chromatin state.^50,51^ However, the integration of our ATAC-seq and scRNA-seq data did not show an enrichment in the expression of genes associated with these open loci in a particular subset, suggesting that HDAC1 does not restrain alternative fates of early-Tex^prog^ cells. Unexpectedly, our analysis revealed that the deletion of HDAC1 led to a closed chromatin state at gene loci in early-Tex^prog^ cells that are later expressed in early-Tex^eff-like^ cells. This indicates that HDAC1 is crucial for establishing an epigenetic potential in early-Tex^prog^ cells that facilitates differentiation towards early-Tex^eff-like^ cells. Our TF-binding motif enrichment analysis revealed that ETS:Runx- and Runx-binding motifs are overrepresented in DARs of WT and HDAC1-cKO early-Tex^prog^ cells, respectively. Since Runx motifs were enriched in DARs of both genotypes, this suggests that Runx family proteins might have different interaction partners and/or are recruited to different genomic loci in early-Tex^prog^ cells dependent on the presence of HDAC1. We therefore propose that HDAC1 regulates the formation and function of a complex TF network required for establishing open chromatin, thereby controlling early-Tex^eff-like^ specification processes via epigenetic processes. However, since HDACs target many non-histone proteins,^52,53^ HDAC1 might also control chromatin accessibility in early-Tex^prog^ cells in a non-epigenetic manner via changing the acetylation status and thus the activity of TFs and other proteins. Finally, given that a fraction of early-Tex^eff-like^ cells 8 days p.i. might directly differentiate from naïve CD8^+^ T cells (i.e. without passing through the early-Tex^prog^ state),^24,35,42^ follow-up studies are required to elucidate trajectories towards early-Tex^eff-like^ cells and the role of HDAC1 during the potential transition of naïve to early-Tex^eff-like^ cells.

CX3CR1^+^ CD8^+^ T cells that emerge at day 8 p.i. substantially contribute to the Tex^eff-like^ cell pool for the following weeks,^22^ In addition, Tex^eff-like^ cells are continuously replenished by Tex^prog^ cells.^20,26,35,42^ This suggests at least two regulatory mechanisms, i.e. self-renewal and replenishment, that control the abundance of Tex^eff-like^ cells once chronic infection is established. By utilizing an inducible deletion system, we demonstrated that HDAC1 is also required for the maintenance of the CX3CR1^+^ Tex cell pool in a CD8^+^ T cell-intrinsic manner. Previous studies have identified several transcriptional and epigenetic regulators controlling Tex^eff-like^ cell differentiation, including T-bet, basic leucine zipper ATF-like transcription factor (BATF), signal transducer and activator of transcription 5A (STAT5A) and SWItch/Sucrose Nonfermenting (SWI/SNF) complexes.^20,22,23,25,26,48,54,55^ In these studies the various regulators were deleted and/or overexpressed before or at the onset of T cell activation, precluding a detailed analysis of whether they are also required for the maintenance of Tex^eff-like^ cells. In contrast, our present study also revealed novel insight into factors controlling the maintenance of Tex^eff-like^ cell subset(s) during chronic infection by showing a key role for HDAC1. It is conceivable that HDAC1 is part of the regulatory program that governs the Tex^eff-like^ cell replenishment process by promoting a Tex^eff-like^ potential in Tex^prog^ cells, similar to its function in early-Tex^prog^ cells. Further studies are required to investigate whether HDAC1 also contributes to the self-renewal process of Tex^eff-like^ cells.

Finally, our results suggest that the modulation of HDAC1 activity and the activation of HDAC1-controlled pathways might serve as an attractive strategy to improve the outcome of T cell-based immunotherapies by promoting the generation of Tex^eff-like^ cells. However, since an excessive effector function of CD8^+^ T cells results in immunopathology,^36,56,57^ caution will be needed to induce an “optimal” degree of effector responses with maximal clinical benefit. Given the role of HDAC1 in a broad range of biological processes,^58^ further studies are necessary to determine whether HDAC1 or HDAC1-regulated pathways are attractive targets for T cell-based immunotherapies against chronic viral infection and potentially cancer.

## Material and Methods

### Mice

Floxed *Hdac1* (MGI:4440556) mice were kindly provided by Patrick Matthias (Friedrich Miescher Institute for Biomedical Research, Basel, Switzerland). *Cd4*-Cre mice (MGI:2386448) were kindly provided by Chris Wilson (University of Washington, Seattle, USA). *Hdac1*^f/f^, *Cd4*-Cre (HDAC1-cKO) mice were previously described^30^ *Rosa26*-STOP-YFP (MGI:2449038) and P14 T cell receptor transgenic (MGI:2665105) mice were kindly provided by Meinrad Busslinger (Research Institute of Molecular Pathology, Vienna, Austria) and Annette Oxenius (ETH Zurich, Zurich, Switzerland), respectively, and were crossed to HDAC1-cKO mice to generate *Rosa26*-STOP-YFP, P14, *Hdac1*^fl/fl^, *Cd4*-Cre (P14-HDAC1-cKO) mice. *Rosa26*-CreERT2 mice (MGI:3699244) were kindly provided by Emilio Casanova (Medical University of Vienna, Vienna, Austria) and were crossed to *Hdac1*^fl/fl^ mice on a *Rosa26*-STOP-YFP background to generate *Rosa26*-STOP-YFP, *Hdac1*^fl/fl^*, Rosa26*-CreERT2 (HDAC1-cKO^CreERT-YFP^). CD45.1 (MGI:4819849) and CD90.1 (MGI:3579311) congenic mice were kindly provided by Jochen Hühn (Helmholtz Center, Braunschweig, Germany) and Annette Oxenius (ETH Zurich, Zurich, Switzerland), respectively. Mice were analyzed at 8-12 weeks of age and were maintained in the Core Facility Laboratory Animal Breeding and Husbandry (CFL) of the Medical University of Vienna. Animal experiments were assessed by the Ethics Committee of the Medical University of Vienna and approved by the Austrian Federal Ministry for Education, Science and Research (animal protocol number: BMBWF-BM BMBWF-66.009/0040-V/3b/2019, BMBWF-66.009/0343-V/3b/2019). Animal husbandry and experiments were performed in compliance with national laws and according to the FELASA (Federation of European Laboratory Animal Science Association) and the ARRIVE (Animal Research: Reporting of In Vivo Experiments) guidelines. Genotyping PCR primers are listed in Table 3.

**Table 3:**
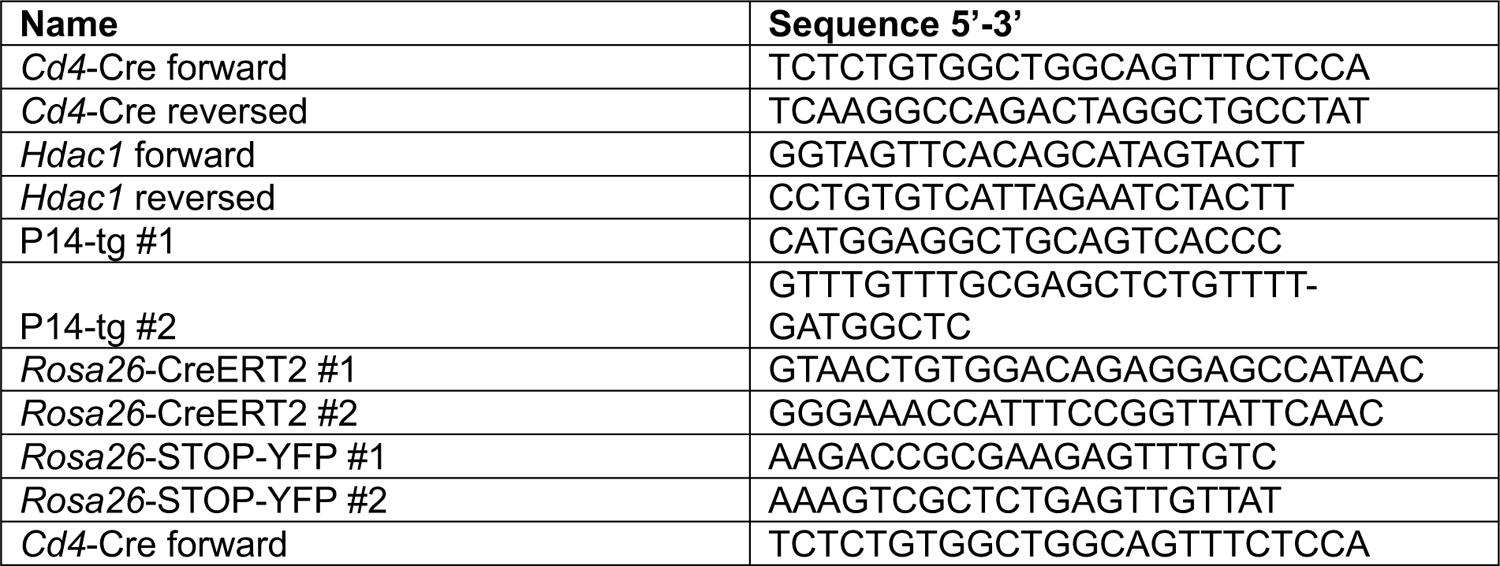
Primer sequences used for genotyping experimental animals.

### LCMV clone 13 (Cl13) infection and measurement of viral titer

LCMV Cl13 was propagated in BHK-21 cells (ATCC No. CCL-10) as described^59,60^ ×10^6^ focusforming units (FFU) of LCMV Cl13. Mice were euthanized 8 or 30 days after infection for further analysis. Viral titers were measured using standard immunofocus assays^61^ with minor modifications. Briefly, Vero cells (ATCC No. CCL-81) were incubated for 1h with blood serum collected from infected mice. Afterwards, the cells were covered for 48h with 1x 199 media (Gibco) containing 0.5% ME agarose (Biozym), 10% FBS (Biowest), 2mM glutamine (Sigma) and 100 U/mL penicillin and streptomycin (Sigma). Cells were fixed with 25% formaldehyde (Sigma) in PBS (Sigma), permeabilized with 1% TRITON X-100/PBS (Promega) and blocked with 5% FBS/PBS. Subsequently, cells were stained with VL-4 rat anti-LCMV nucleoprotein (BioXCell), followed by staining with anti-rat horseradish peroxidase (Jackson Immunoresearch) in 2% FBS/PBS. Infected cells were visualized using AEC chromogen and substrate (Biolegend). Viral titers were calculated based on the number of colored “spots” on the plate and the dilution factor of the serum samples.

### Generation of mixed bone marrow (BM) chimeric mice

BM cells isolated from either WT or HDAC1-cKO mice (both expressing the congenic marker CD45.2) were mixed at a 1:1 ratio with BM cells from WT CD45.1 mice. Mixed BM cells (2×10^6^) were transferred i.v. into lethally irradiated CD45.1^+^ mice. Reconstituted mice were i.v. infected with 2×10^6^ FFU of LCMV Cl13 7-8 weeks after transplantation. Eight days later, mice were euthanized and splenocytes were analyzed by flow cytometry.

### CD8^+^ T cell transfer followed by LCMV Cl13 infection

Naïve P14-WT and P14-HDAC1-cKO CD8^+^ T cells (CD90.2^+^) were isolated as described previously^62^ and mixed at a 1:1 ratio. Either 1× 10^6^ (analyzed at 48 and 67 hours post infection) or 2× 10^4^ (analyzed on 5 and 8 days post infection) of the mixed P14 CD8^+^ T cells were transferred i.v. into CD90.1^+^ congenic mice. One day after the transfer, recipient mice were infected i.v. with 2×10^6^ FFU of LCMV Cl13. Mice were euthanized at various time-points after infection as indicated and splenocytes were analyzed using flow cytometry.

### Tamoxifen treatment

For timed induction of *Hdac1* deletion, HDAC1-cKO^CreERT-YFP^ mice were treated with 2mg of Tamoxifen (Sigma) intraperitoneally (i.p.) after chronic disease has been established (i.e. 9 days post infection). 6 days later (i.e. 15 days post infection), mice were euthanized and splenocytes were analyzed by flow cytometry.

### Flow cytometry analysis

Single cell suspensions were prepared by mashing organs through a 70 μm cell strainer (Corning or pluriSelect). Hepatic lymphocytes were enriched via 35% Percoll (Cytiva)/RPMI 1640 (Sigma) gradient centrifugation (700× g for 20min at RT, no breaks). Erythrocytes were lysed using 1× BD Pharm Lyse Buffer according to the manufacturer’s instruction (BD Biosciences). Cells were incubated with Fc block (1:250; BD Biosciences) and Fixable Viability Dye eFluor 506 (1:1000; Thermo Fisher Scientific). For the detection of viral-specific CD8^+^ T cells, cells were subsequently incubated with fluorescently labeled H2-D^b^/GP33-41 tetramer (1:1000; kindly provided by the NIH Tetramer Core Facility), followed by staining with antibodies for surface markers. Intracellular stainings were performed using the Foxp3/Transcription Factor Staining Buffer Set (Thermo Fischer Scientific) according to the manufacturer’s instruction. For intra-cytoplasmic detection of active-caspase 3, splenocyte were incubated at 37°C for 5h in complete RPMI1640 medium, supplemented with 10% FBS, 100 U/ml penicillin-streptomycin (Thermo Fisher Scientific), 2 mM L-glutamin (Thermo Fisher Scientific), 0.1 mM non-essential amino acid (Thermo Fisher Scientific), 1 mM sodium pyruvate (Sigma), 55 μM of β-mercaptoethanol (Sigma). Subsequently, cells were fixed with BD Cytofix Fixation Buffer and were stained with PE Rabbit Anti-Active Caspase-3 (BD Biosciences) in BD Perm/Wash Buffer (BD Biosciences) according to the manufacturer’s instructions.^20^ Cells were acquired with a BD LSRFortessa (BD Biosciences) or CytoFLEX (Beckman Coulter) and analyzed using FlowJo v10.8.1 software (BD Biosciences). Antibodies used in this study are listed in Table 4.

**Table 4:** Antibody list.

| Epitope | Clone | Vendor |
| --- | --- | --- |
| anti-GFP (biniding to YFP) | 1A12-6-18 | BD Biosciences |
| anti-mouse CD107a | 1D4B | Biolegend |
| anti-mouse CD107b | M3/84 | Biolegend |
| anti-mouse CD45.1 | A20 | Biolegend |
| anti-mouse CD45.2 | 104 | Biolegend |
| anti-mouse CD69 | H1.2F3 | Thermo Fisher Scientific |
| anti-mouse CD8α | 53-6.7 | Biolegend |
| anti-mouse CD90.2 | 53-2.1 | Biolegend |
| anti-mouse CX3CR1 | SA011F11 | Biolegend |
| anti-mouse Eomes | Dan11mag | Thermo Fisher Scientific |
| anti-mouse HDAC1 | Sat13 | kindly provided by<br>Christian Seiser |
| anti-mouse IFN-γ | XMG1.2 | Biolegend |
| Anti-mouse Ki67 | 16A8 | Biolegend |
| anti-mouse Ly108 | 330-AJ | Biolegend |
| anti-mouse PD1 | 29F.1A12 | Biolegend |
| anti-mouse Perforin | s16009a | Biolegend |
| anti-mouse TCR Vα2 | B20.1 | Biolegend |
| anti-mouse Tim3 | RMT3-23 | Biolegend |
| anti-mouse TNF-α | MP6-XT22 | Biolegend |
| anti-mouse/human Active Caspase-3 | C92-605.rMAb | BD Biosciences |
| anti-mouse/human CD44 | IM7 | Biolegend |
| anti-mouse/human KLRG1 | 2F1/KLRG1 | Biolegend |
| anti-mouse/human T-bet | 4B10 | Biolegend |
| anti-mouse/human TCF-7/ TCF-1 | S33-966 | BD Biosciences |
| anti-rat CD90/mouse CD90.1 | OX-7 | Biolegend |
| goat anti-rabbit IgG Alexa Fluor 467 |  | Abcam |
| goat anti-rabbit IgG Alexa Fluor 488 |  | Thermo Fisher Scientific |
| H2-D<sup>b</sup>/GP33-41 tetramer |  | kindly provided by<br>NIH Tetramer Core Facility |
| TotalSeq™-A0301 anti-mouse Hashtag 1 | (M1/42; 30-F11) | Biolegend |
| TotalSeq™-A0302 anti-mouse Hashtag 2 | (M1/42; 30-F11) | Biolegend |
| TotalSeq™-A0303 anti-mouse Hashtag 3 | (M1/42; 30-F11) | Biolegend |
| TotalSeq™-A0303 anti-mouse Hashtag 3 | (M1/42; 30-F11) | Biolegend |

### Dimensionality reduction of flow cytometry data

Dimensionality reduction was performed using the DownSample and t-SNE Plugins in FlowJo™ v10.8.1 (BD Biosciences). For the generation of t-SNE plots, WT and HDAC1-cKO biological replicates were down-sampled to the same number of cells/sample and concatenated for further analysis. t-SNE was run on the concatenated file using the default parameters provided by the software.

### Single-cell RNA sequencing (scRNA-seq) of naïve and virus-specific CD8^+^ T cells

Single cell suspensions from the spleen of either LCMV-infected (infected as described above) or uninfected WT and HDAC1-cKO mice were prepared. Equal numbers of splenocytes from one female and one male mouse were combined for each scRNA-seq sample per condition. Samples were incubated with fluorescently labeled H2-D^b^/GP33-41 tetramer, appropriate antibodies against surface markers, oligonucleotide-conjugated antibodies (TotalSeq-A hashtags; Biolegend) and fixable viability dye (Zombie Green; Biolegend). Viable naïve (CD90.2^+^CD8α^+^CD44^-^) and LCMV-specific (CD90.2^+^CD8α^+^CD44^+^GP33-tet^+^) WT and HDAC1-cKO CD8^+^ T cells were sorted using a SH800S Cell Sorter (Sony Biotechnology). These four samples were pooled for multiplex sequencing (approx. 30,000 cells/run; a total of two runs was performed). Libraries were generated using the Chromium Controller and the Next GEM Single Cell 3’ Reagent Kit (v3.1, 10x Genomics) according to the manufacturer’s instructions. Libraries were sequenced by the Biomedical Sequencing Facility at CeMM using the Illumina NovaSeq 6000 platform. Raw sequencing results were converted to gene-barcode-count matrices using Cell Ranger (v6.1.2, 10x Genomics). The raw count matrices, which recovered 11,813 (1^st^ run) and 18,363 (2^nd^ run) cells with 32,289 genes detected from both replicates respectively, have been used for further computational analysis.

### Computational analysis of scRNA-seq data

Analysis was performed in R (version 4.1.2) using the Seurat package (version 4.2.0) as well as tidyverse (version 2.0.0), ggplot2 (version 3.4.2) and EnhancedVolcano (version 1.18.0). Replicates have been demultiplexed with a positive quantile threshold of 0.99 (HTOdemux() function) and resulted in a total of 7,108 Negative, 2,398 Doublet and 20,670 Singlet cells. Only Singlets were used for cell type annotation with SingleR (version 1.8.1). All cells annotated as “T cells” or “NK cells” were kept (20,171 cells) for further analysis. The number of genes (nGene), number of UMIs (nUMI), percentage of mitochondrial genes (percent.mt) and the novelty score (number of genes detected per UMI (log10GenesPerUMI)) were used for quality control. Following filters were applied: 300 ≤ nGene ≤ 3000, nUMI ≥ 500, percent.mt ≤ 5 and log10GenesPerUMI > 0.8. Before integrating the data from the two replicates, the 16,393 high-quality cells were used to regress out unwanted variations due to the expression of cell cycle genes (SCTransform() and CellCycleScoring()). Clustering was performed at different resolutions considering all principal components (PCs) for which the change of variation to the subsequent PC was larger than 0.1%. A clustering resolution of 0.6 was used for all following analyses and visualizations. Marker genes for all clusters were defined using FindAllMarkers() with the default settings. AddModuleScore() followed by FeaturePlot() was used to visualize the expression of previously published genes. Differentially expressed genes (DEGs) between groups were determined with FindMarkers() where the minimum log_2_FC was set to 0 and the minimum fraction of cells in which a gene has to be detected in either of the two groups to be tested was set to 1%. Gene Ontology (GO) term enrichment analyses were conducted with the packages clusterProfiler (version 4.8.1), org.Mm.eg.db (version 3.17.0) and AnnotationHub (version 3.8.0). Top 200 marker genes of each cluster were used and resulting top 5 enriched pathways (in Biological Processes) from each cluster were collapsed to parent terms using Revigo (version 1.8.1) and then visualized with dotplot() from the enrichplot package (version 1.20.0).

### Sample preparation of virus-specific CD8^+^ T cells for ATAC-seq analysis

Naïve WT and HDAC1-cKO P14 T cells (CD90.2^+^) were isolated as described previously^62^ and 1×10^5^ cells were transferred i.v. into CD90.1^+^ congenic mice. One day after the transfer, recipient mice were infected i.v. with 2×10^6^ FFU of LCMV Cl13. Eight days later, mice were euthanized and splenocytes from 4-5 female mice were pooled for staining and subsequent cell sorting. 5×10^4^ early-Tex^prog^ (CD90.2^+^CD8α^+^Tim3^-^Ly108^+^) and 5×10^4^ early-non-Tex^prog^ (CD90.2^+^CD8α^+^Tim3^+^Ly108^-^) cells were sorted into 1× PBS. High-throughput chromatin accessibility mapping (ATAC-seq) was performed as previously described,^63,64^ with minor modifications. After centrifugation at 500× g for 5min at 4°C, the cell pellets were resuspended in tagmentation-mix (containing Tagment DNA buffer, Tagment DNA Enzyme (Illumina), Digitonin (Promega) and Proteinase K (Roche)) and incubated for 30min at 37°C with shaking (300rpm). The reaction was stopped by incubating the samples on ice. For DNA isolation, the MinElute PCR Purification Kit (Qiagen) was used according to the manufacturer’s instructions and eluted into 12 μl of elution buffer. 1 µl eluate was used in a quantitative PCR (qPCR) reaction to estimate the optimal number of cycles for library amplification. The remaining sample of each tagmented sample was then amplified corresponding to the C_q_ value (i.e., the cycle number at which fluorescence has increased above background levels) in the presence of custom Nextera primers. PCR amplification was followed by SPRI (Beckman Coulter) size selection to exclude fragments larger than 1,200 bp. The DNA concertation of each library was assessed with a Qubit 2.0 Fluorometric Quantitation system (Life Technologies, Carlsbad, CA, USA), before pooling in equimolar amounts. The resulting pools were sequenced on a NovaSeq 6000 instrument (Illumina, San Diego, CA, USA) in 50 base pair paired-end configuration. Chromatin accessibility mapping by ATAC-seq was done in [three] biological replicates. NGS reads in unaligned BAM files were converted into FASTQ format with samtools, NGS adapter sequences were removed via fastp (0.23.2, GTCTCGTGGGCTCGG) and the reads were aligned to the GRCm38 (UCSC Genome Browser mm10) assembly with Bowtie2 (2.4.5, --very-sensitive, --no-discordant, --maxins 2000), before deduplicating with samblaster (0.1.24). Alignment statistics and mitochondrial read fractions were collected with samtools. Peaks were called with MACS2 (2.2.7.1, --no-model, --keep-dup auto, --extsize 147) and both, peak annotation and motifs discovery was performed with HOMER (4.11). UCSC Track hubs allowed for visual inspection of results. A consensus peak set was compiled and all peaks in mm10 assembly regions blacklisted by the ENCODE project^65^ were removed. Sequencing and mapping analysis were performed by the Biomedical Sequencing Facility at CeMM. Resulting count matrices and peak files were used for further computational analysis.

### Computational analysis of ATAC-seq data

Differentially accessible regions (DARs) in early-Tex^prog^ and early-non-Tex^prog^ were identified with the R (version 4.1.2) package DESeq2 (version 1.40.2) using the default log_2_FC threshold and an alpha of 0.05. Genes associated to DARs were used to calculate module scores (AddModuleScore() from the Seurat package (version 4.2.0)) to visualize the respective gene expression in the scRNA-seq clusters. For statistical testing an ANOVA followed by Tukey’s test was performed. The same set of genes was also used to conduct the GO term enrichment analyses for Biological Processes with the same R packages as described above. Top 10 enriched pathways from each set were collapsed to parent terms using Revigo (version 1.8.1), and then visualized with dotplot() from the enrichplot package (version 1.20.0). Gene Set Enrichment Analysis (GSEA) against a known Effector Signature (GSE9650_EFFECTOR_VS_EXHAUSTED_CD8_TCELL_UP) from the ImmuneSigDB were carried out using the packages msigdbr (version 7.5.1) and fgsea (version 1.26.0). Transcription factor motifs in DARs were determined by de novo motif analysis using findMotifsGenome.pl from HOMER (v4.10.3). Parameters were set to mask repeat regions (-mask), use the exact input region sizes (-size given) and find maximum ten motifs per length (-S 10). *p*-values < 10^-11^ were considered true positive.

### Data and code availability

Raw scRNA-seq and ATAC-seq data (sequence, count and/or peak data) will become available upon acceptance for publication. Used scripts and relevant metadata are available on GitHub (github.com/medunivienna-IFI-immunobiology/2024_Rica_HDAC1-CD8Tcells-chronicLCMV).

### Statistical Analysis

Statistical analysis was performed using Prism 8 or 9 (GraphPad). The values of the geometric mean fluorescence intensity (gMFI) were normalized by dividing the value of each sample by the average of the corresponding WT samples. Unpaired or paired two-tailed Student’s *t*-test was used for comparisons between two groups. One-way ANOVA followed by Tukeýs multiple-comparison test was used for the comparison of more than 2 groups. The *p*-values are defined as follows: *, *p* <0.05; **, *p* <0.01; ***, *p* <0.001; n.s., not significant.

## Acknowledgments

We thank the NIH Tetramer Facility (Atlanta, GA) for providing MHC class I tetramers, Ms. Kirkley and Mr. Klimek (CeMM, Vienna, Austria) for their assistance with sorting of cells isolated from LCMV-infected mice, and the MedUni Wien core facility laboratory animal breeding and husbandry (CFL) for animal husbandry. Figure graphics were created with BioRender.com. S.S. was supported by Austrian Science Fund (FWF) projects: P27747 and P35436. W.E. was supported by Austrian Science Fund (FWF) projects: F7001, F7005, P35372; and by the FWF and Medical University of Vienna doctoral programs (DK W1212) “Inflammation and Immunity” and (DOC 32 doc.fund) “TissueHome”. N.B. was supported by Austrian Science Fund (FWF) projects: F7004 and P30885. C.B. was supported by Austrian Science Fund (FWF) project: F7002.

## Author contributions

Conceptualization, W.E. and S.S.; Methodology, R.R., T.K. and S.S.; Formal Analysis, R.R., M.W., M.S., and S.S.; Investigation, R.R., M.S., E.M., L.S., V.S., D.W. and S.S.; Resources, C.B.; Data Curation, M.W.; Writing – Original Draft, R.R., M.W., W.E., and S.S.; Visualization, R.R., M.W. and S.S.; Supervision, N.B., W.E. and S.S.; Funding Acquisition, W.E. and S.S.

---

**Fig. S1:**
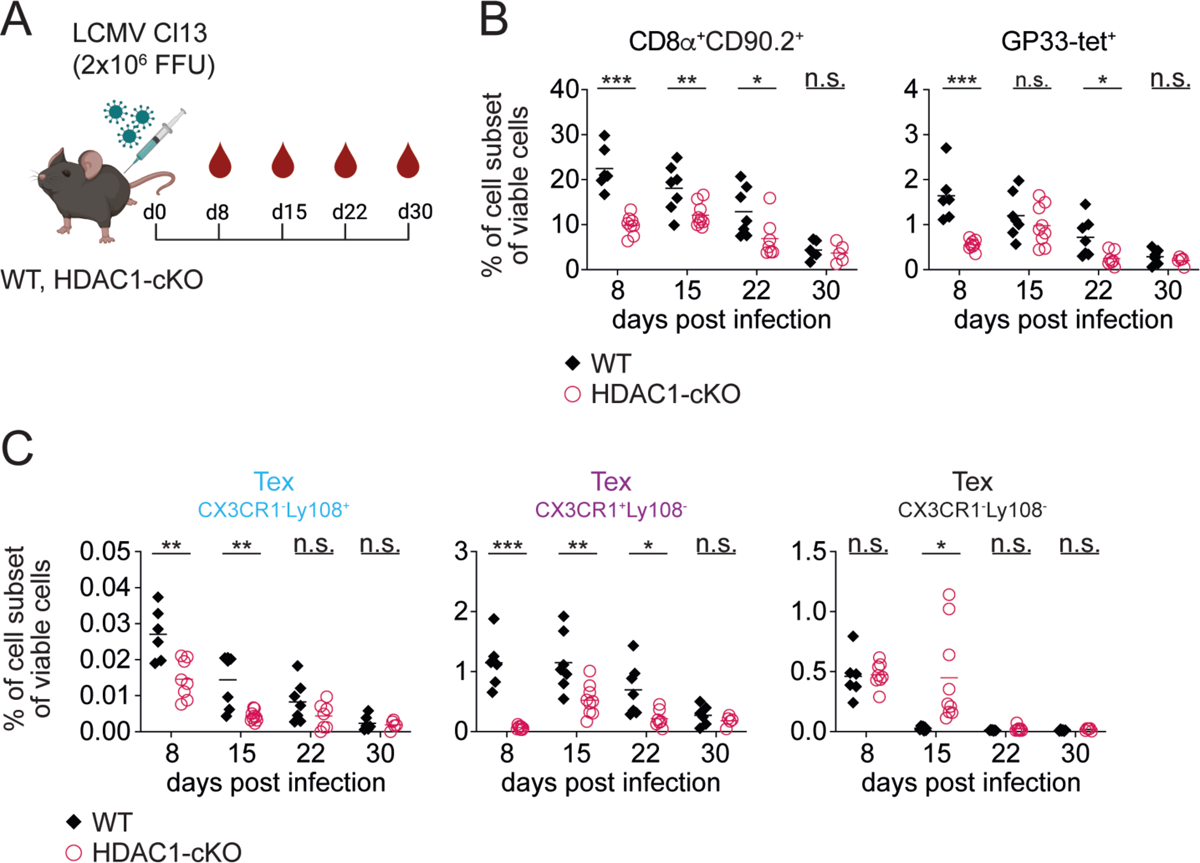
Differentiation kinetics of circulating Tex cell subsets during the course of chronic infection. (A) Scheme of the experimental design. WT and HDAC1-cKO mice were infected with LCMV Cl13. On day 8 (d8), 15, 22 and 30 post infection (p.i.) blood was drained to assess Tex cell subsets. (B) Diagrams show the summary of the percentage of CD8^+^ T cells (CD8α^+^CD90.2^+^) and Tex cells (GP33-tet^+^, pre-gated on CD8α^+^CD90.2^+^) of viable circulating lymphocytes. (C) Summary diagrams show percentages of Tex cell subsets: CX3CR1^-^Ly108^+^ Tex (left), CX3CR1^+^Ly108^-^ Tex (middle) and CX3CR1^-^Ly108^-^ Tex (right) cells of viable circulating lymphocytes. (B,C) Diagrams show the summary of 5-9 mice per group analyzed in 2 independent experiments. Horizontal bars indicate the mean. \**p* < 0.05, \*\**p* < 0.01, \*\*\**p* < 0.001, n.s. *p* ≥ 0.05. An unpaired two-tailed Student’s *t*-test was used to compare WT vs HDAC1-cKOcells on the indicated day.

**Fig. S2:**
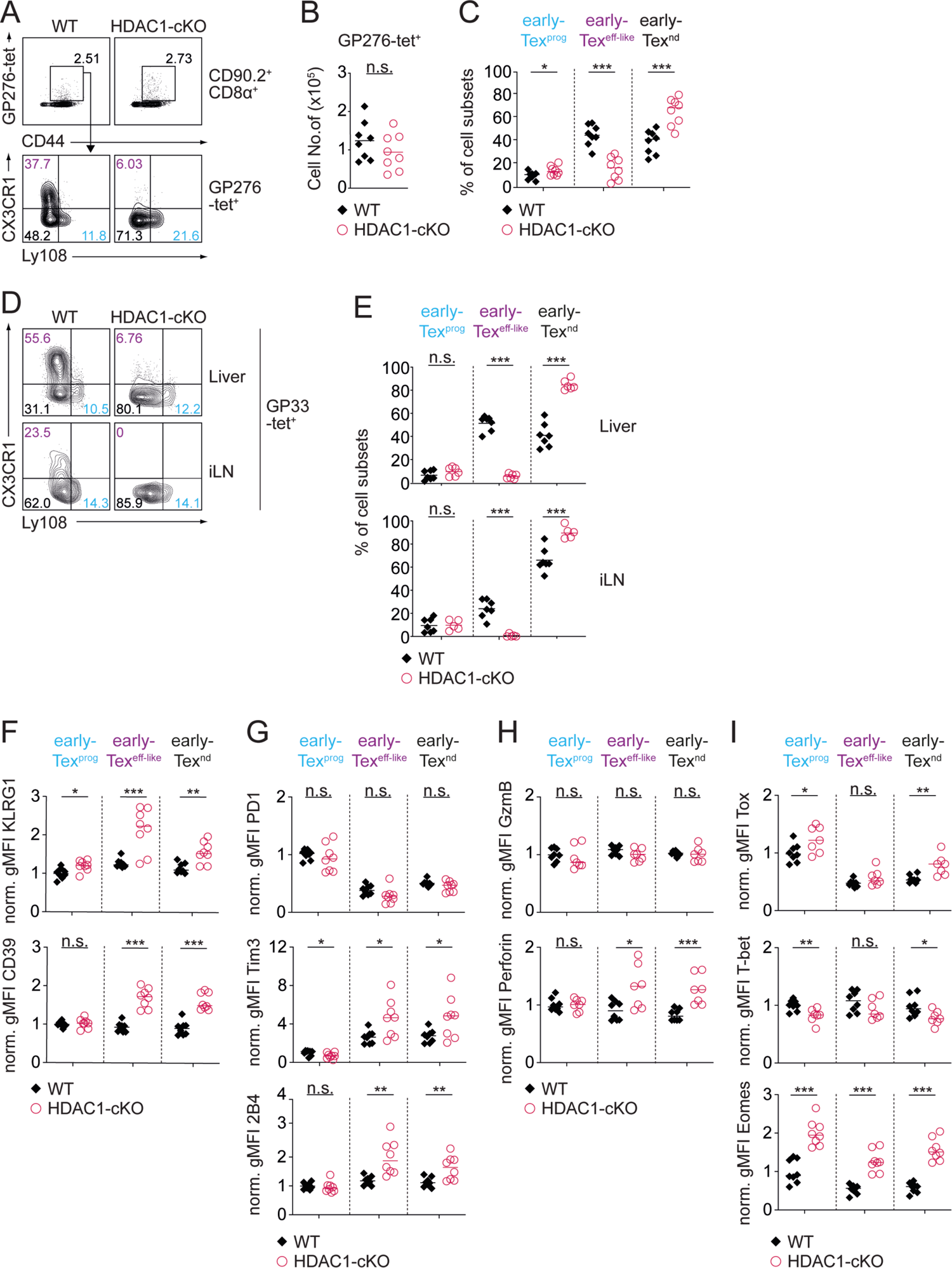
Altered Tex cell subset composition is independent of the TCR-specificity and occurs systemically. (A) Contour plots show GP276-tet^+^ staining and CD44 expression (upper panel) and the distribution of Tex cell subsets in WT and HDAC1-cKO mice on d8 post infection (p.i.): early-Tex^prog^ (CX3CR1^-^Ly108^+^) (blue), early-Tex^eff-like^ (CX3CR1^+^Ly108^-^) (purple), and early-Tex^nd^ (CX3CR1^-^Ly108^-^) (black) (lower panel). (B) Summary diagram shows absolute numbers of GP276-tet^+^ Tex cells. (C) Summary diagram depicts percentages of Tex cell subsets as shown in (A). (D) Contour plots show distribution of GP33-tet^+^ Tex cell subsets in liver and inguinal lymph nodes (iLN) on d8 p.i. (E) Diagrams show summary of the percentages of Tex cell subsets as depicted in (D). (F) Diagrams display normalized gMFI (geometric mean fluorescence intensity; average WT early-Tex^prog^ expression set to 1) expression of KLRG1 (upper panel) or CD39 (lower panel) in all Tex cell subsets. (G) Summary diagrams show normalized gMFI expression of PD1 (upper panel), Tim3 (middle panel) or 2B4 (lower panel) in all Tex cell subsets. (H) Diagrams show normalized gMFI expression of granzyme B (GzmB) (upper panel), and Perforin (lower panel) in all Tex cell subsets. (I) Summary of normalized gMFI expression of the transcription factors Tox (upper panel), T-bet (middle panel) and Eomes (lower panel) in all Tex cell subsets. Numbers in the plots (A,D) depicts the percentage of cells within the indicated regions. (F-I) Cells were pre-gated on CD90.2^+^CD8α^+^ GP33-tet^+^CD44^+^. Data are representative (A,D) or show the summary (B,C,E-I) of 8 mice (B,C,F-I), 5-7 mice (E) per genotype analyzed in 2 independent experiments. \**p* < 0.05, \*\**p* < 0.01, \*\*\**p* < 0.001, n.s. *p* ≥ 0.05. An unpaired two-tailed Student’s *t*-test was used to compare WT vs HDAC1-cKO cells.

**Fig. S3:**
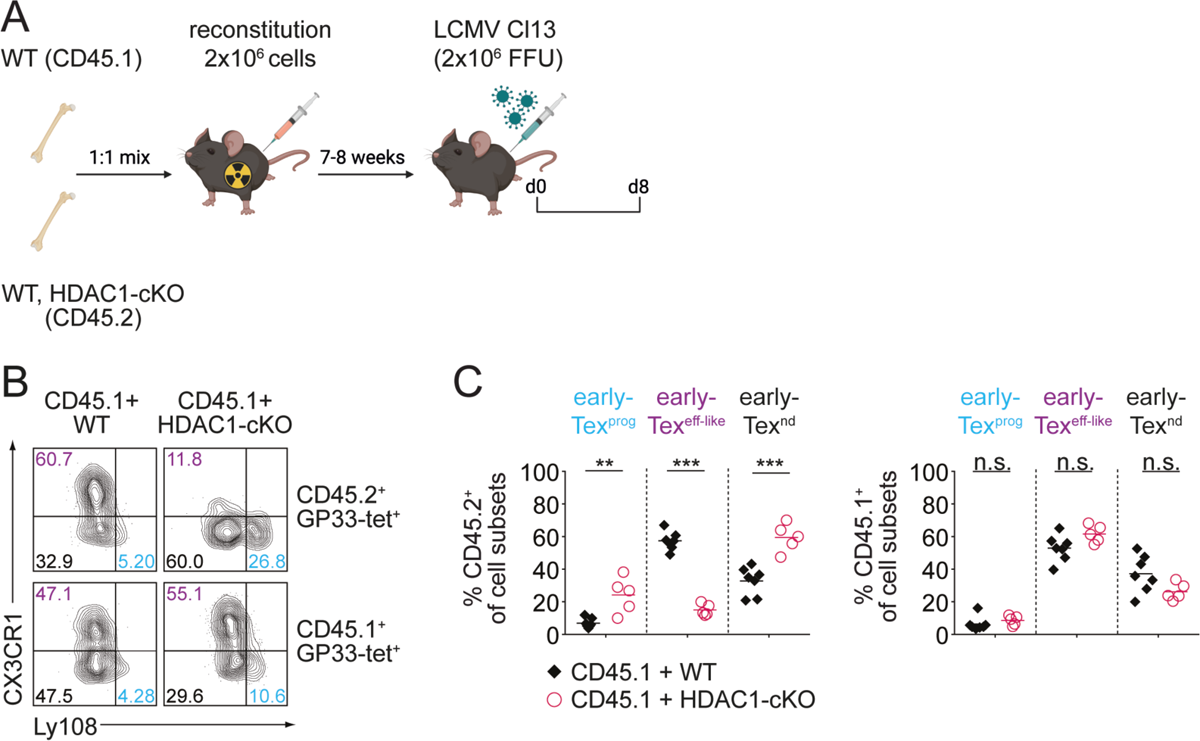
HDAC1 controls early-Tex^eff-like^ subset differentiation in a CD8^+^ T cell-intrinsic manner. (A) Schematic drawing of the experimental design. Bone marrow (BM) from either WT (CD45.2^+^) or HDAC1-cKO (CD45.2^+^) mice was mixed with BM from WT CD45.1^+^ mice at a 1:1 ratio and transferred into irradiated CD45.1^+^ recipient mice. Reconstituted mice were infected with LCMV Cl13 7-8 weeks after BM transplantation. Eight days later (d8) splenocytes were analyzed by flow cytometry. (B) Contour plots show the distribution of Tex cell subsets on d8 p.i.: early-Tex^prog^ (CX3CR1^-^Ly108^+^) (blue), early-Tex^eff-like^ (CX3CR1^+^Ly108^-^) (purple) and early-Tex^nd^ (CX3CR1^-^Ly108^-^) (black) within WT + CD45.1 or HDAC1-cKO + CD45.1 recipient mice. Cells were pre-gated on CD90.2^+^CD8α^+^GP33-tet^+^CD44^+^ cells. WT and HDAC1-cKO (CD45.2^+^) cells are shown in the upper panel, whereas the control WT (CD45.1^+^) cells are shown in the lower panel. (C) Summary diagrams show the percentages of Tex cell subsets within the CD45.2^+^ (left) and the CD45.1^+^ (right) cell compartments. Numbers in the plots (B) show the percentage of cells within the indicated regions. Data are representative (B) or show the summary (C) of 5-7 mice per genotype analyzed in 2 independent experiments. (C) Horizontal bars indicate the mean. \**p* < 0.05, \*\**p* < 0.01, \*\*\**p* < 0.001, n.s. *p* ≥ 0.05. An unpaired two-tailed Student’s *t*-test was used to compare WT vs HDAC1-cKO cells.

**Fig. S4:**
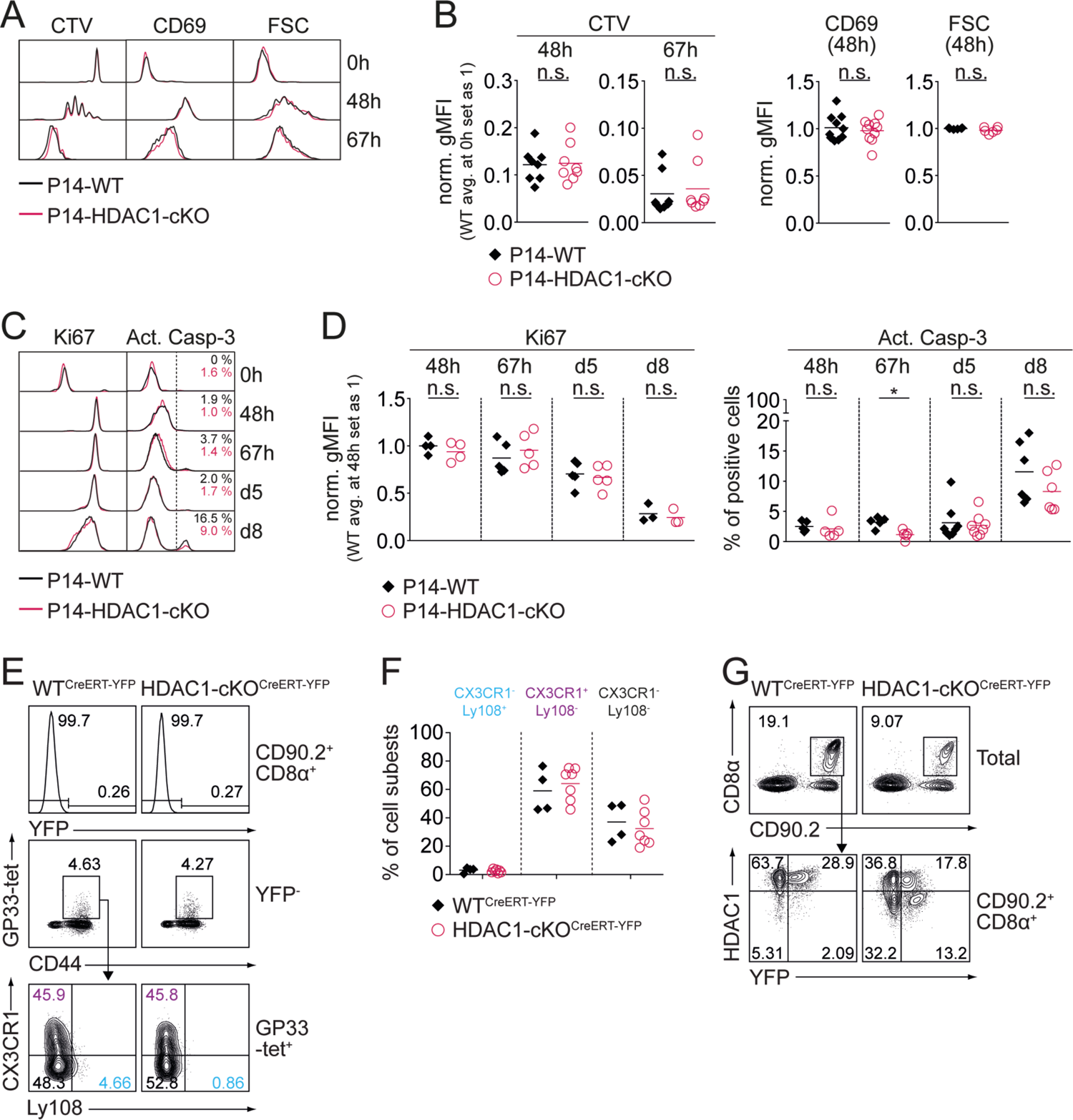
HDAC1 controls early-Tex^eff-like^ subset differentiation and maintenance. (A) Histograms show cell proliferation (Cell Trave Violet – CTV), activation marker CD69 expression and forward scatter (FSC) of P14-WT and P14-HDAC1-cKO CD8^+^ T cells at 48h and 67h post infection (p.i.). (B) Diagrams show relative expression levels (gMFI) of markers depicted in (A). Expression levels in P14-WT^R26-YFP^ cells at timepoint 0h (for CTV) or 48h (for CD69 and FSC) were set as 1. (C) Histograms show the expression of the proliferation marker Ki67 and the apoptosis marker active Caspase-3 (Act. Casp-3) in P14-WT and P14-HDAC1-cKO cells CD8^+^ T cells at the indicated time points of infection (0h, 48h, 67h, d5 and d8). (D) Diagrams show relative expression levels (gMFI) of Ki67 (left) and the percentages of cells that are active Casp-3 positive (right) as depicted in (C). (E) Upper panel: histograms show percentage of YFP-expressing WT^CreERT-YFP^ and HDAC1-cKO^CreERT-YFP^ CD8^+^ T cells (CD90.2^+^CD8α^+^) on day8 (d8) post infection (p.i.). Middle panel: YFP^-^ cells were further gated on GP33-tet^+^ Tex cells. Lower panel: Distribution of Tex cell subsets is shown: CX3CR1^-^Ly108^+^ Tex (blue), CX3CR1^+^Ly108^-^Tex (purple) and CX3CR1^-^Ly108^-^ Tex (black). (F) Summary diagrams show percentages of Tex cell subsets as depicted in (E). (G) Contour plots show HDAC1 and YFP co-expression in WT^CreERT-YFP^ and HDAC1-cKO^CreERT-YFP^ CD8^+^ T cells (CD90.2^+^CD8α^+^) after tamoxifen administration on d15 p.i. Numbers in the histograms (C) or plots (E,G) represent the percentage of cells in the indicated regions. Data are representative (A,C,E,G) or show the summary (B,D) of 6-10 (B) or 3-8 (D) mice per genotype analyzed in 3-5 (B) and 2-4 (D) independent experiments or (F) of 4-6 mice per genotype from 2 independent experiments. (B,D,F) Horizontal bars indicate the mean. \**p* < 0.05, \*\**p* < 0.01, \*\*\**p* < 0.001, n.s. *p* ≥ 0.05. An unpaired two-tailed Student’s *t*-test was used to compare P14-WT vs P14-HDAC1-cKO cells or WT^CreERT-YFP^ vs HDAC1-cKO^CreERT-YFP^ cells.

**Fig. S5:**
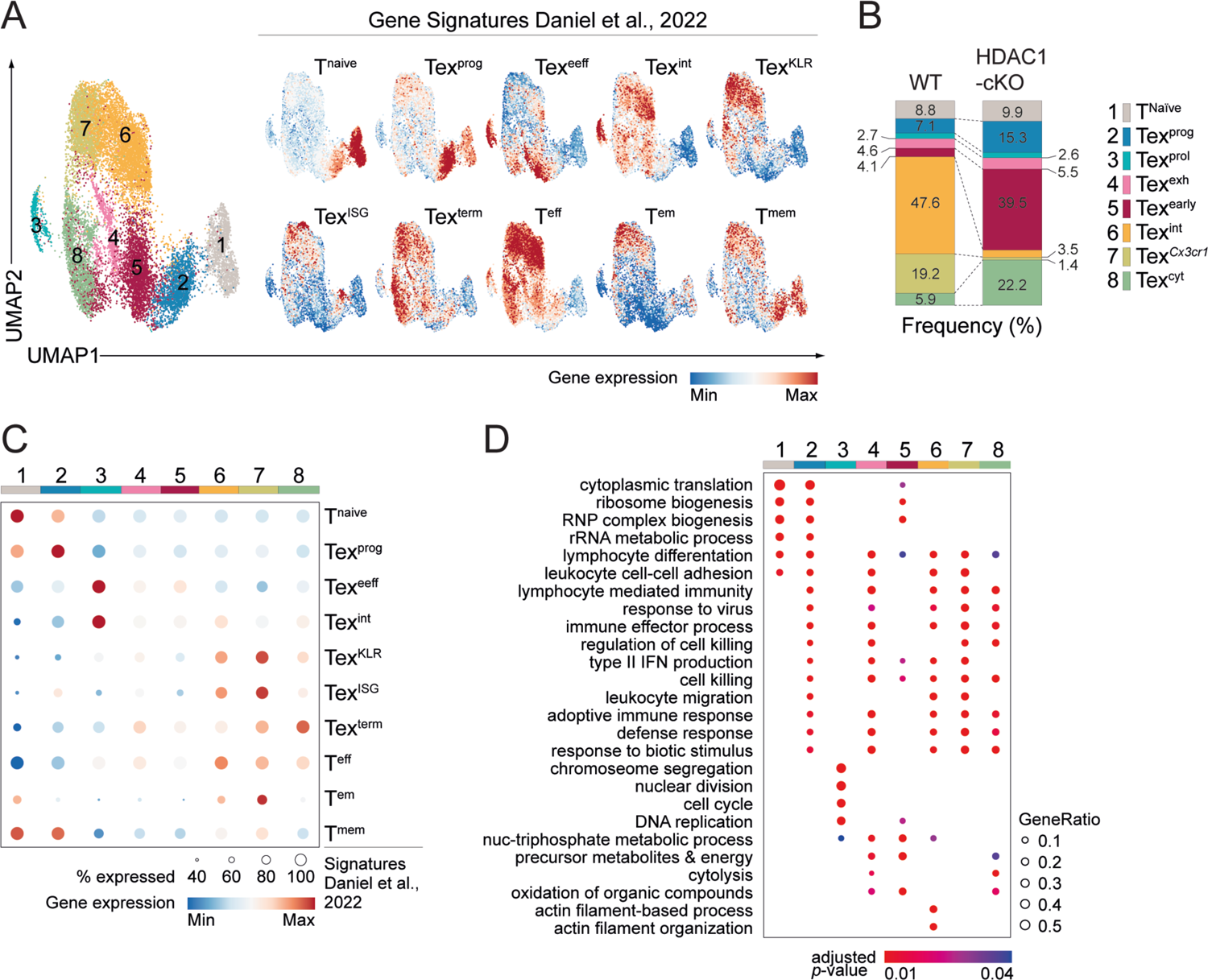
scRNA-seq reveals transcriptionally distinct cell clusters between LCMV-specific WT and HDAC1-deficient CD8^+^ T cells. (A) UMAPs show clustering of naïve and virusspecific WT and HDAC1-cKO CD8^+^ T cells from non-infected or day 8 LCMV Cl13-infected mice (left). Feature plots (right) show the expression of gene signatures from a published data set (Daniel et al., 2022) projected onto our scRNA-seq data. Contrast was improved by using minimum and/or maximum cut-offs and positive cells were plotted on top. (B) Stacked bar graph depicts the frequency of each cluster within the total of WT and HDAC1-cKO CD8^+^ T cells. (C) Bubble plot shows comparison of gene signatures of WT and HDAC1-cKO clusters to a published dataset (Daniel et al., 2022). Bubble size corresponds to the percentage of genes overlapping with the published gene sets, bubble color to the average expression of overlapping genes. (D) Bubble plot showing significantly (adjusted *p*-value < 0.05) enriched GO terms (biological processes) in all clusters obtained from clusterProfiler. Top 200 markers of each cluster were used for enrichment analysis. Top 5 enriched pathways from each cluster were collapsed to parent terms using Revigo and then visualized. Bubble size corresponds to the ratio of marker genes overlapping with the given pathway (Gene Ratio), bubble color corresponds to the adjusted *p*-value. Numbers on plots represent the cluster number as assigned by scRNA-seq analysis (A) or the frequencies within the indicated regions (B).

**Fig. S6:**
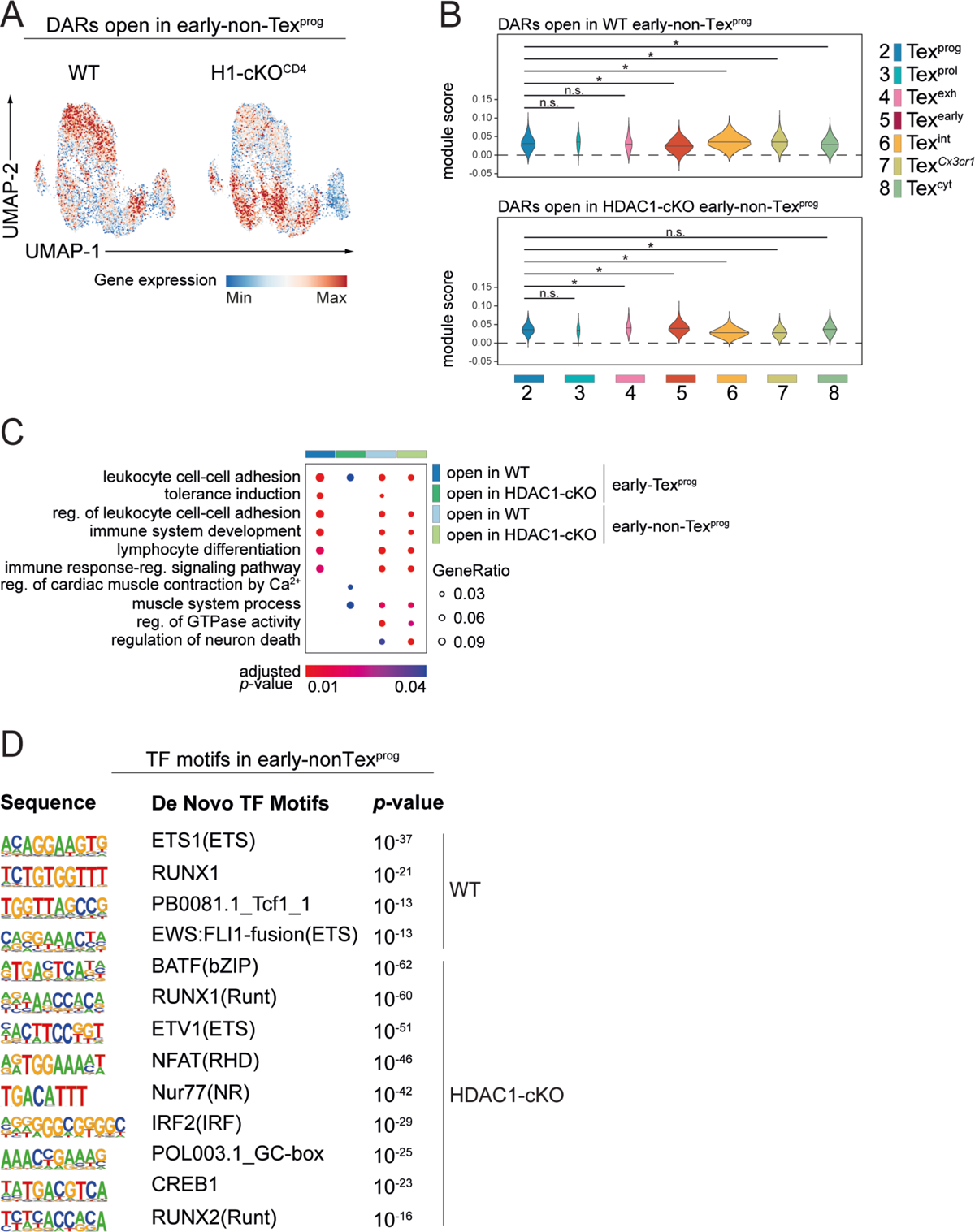
HDAC1 alters the chromatin landscape required for the maintenance of the Tex-^eff-like^ cell pool. (A) Expression of gene transcripts in scRNA-seq of genes associated with open DARs in early-non-Tex^prog^ from ATAC-seq. (B) Violin plots depict the average expression of genes associated with open DARs in WT early-nonTex^prog^ cells (upper panel), or HDAC1-cKO early-non-Tex^prog^ cells (lower panel), calculated as module scores for clusters C2-C8 as defined by scRNA-seq. ANOVA followed by Tukey’s test was performed, * adjusted *p*-value < 0.05, n.s. not significant. (C) Bubble plot showing significantly enriched (adjusted *p*-value < 0.05) GO terms (biological processes) in genes associated with open DARs of WT and HDAC1-cKO early-Tex^prog^ and early-non-Tex^prog^ cells. Bubble size corresponds to the ratio of genes overlapping with the given pathway (Gene Ratio), bubble color corresponds to the adjusted *p*-value. (D) Transcription factor (TF) motifs accessible in WT and HDCA1-cKO early-non-Tex^prog^ cells as determined by de novo motif analysis using HOMER (*p*-values < 10^-11^ were considered true positive).

## References

1. Blank, C.U., Haining, W.N., Held, W., Hogan, P.G., Kallies, A., Lugli, E., Lynn, R.C., Philip, M., Rao, A., Restifo, N.P., et al. (2019). Defining ‘T cell exhaustion’. Nat Rev Immunol 19, 665–674. 10.1038/s41577-019-0221-9.

2. McLane, L.M., Abdel-Hakeem, M.S., and Wherry, E.J. (2019). CD8 T Cell Exhaustion During Chronic Viral Infection and Cancer. Annu Rev Immunol 37, 457–495. 10.1146/annurev-immunol-041015-055318.

3. Kallies, A., Zehn, D., and Utzschneider, D.T. (2020). Precursor exhausted T cells: key to successful immunotherapy? Nat Rev Immunol 20, 128–136. 10.1038/s41577-019-0223-7.

4. Speiser, D.E., Utzschneider, D.T., Oberle, S.G., Munz, C., Romero, P., and Zehn, D. (2014). T cell differentiation in chronic infection and cancer: functional adaptation or exhaustion? Nat Rev Immunol 14, 768–774. 10.1038/nri3740.

5. Jin, X., Bauer, D.E., Tuttleton, S.E., Lewin, S., Gettie, A., Blanchard, J., Irwin, C.E., Safrit, J.T., Mittler, J., Weinberger, L., et al. (1999). Dramatic rise in plasma viremia after CD8(+) T cell depletion in simian immunodeficiency virus-infected macaques. J Exp Med 189, 991–998. 10.1084/jem.189.6.991.

6. Johnston, R.J., Comps-Agrar, L., Hackney, J., Yu, X., Huseni, M., Yang, Y., Park, S., Javinal, V., Chiu, H., Irving, B., et al. (2014). The immunoreceptor TIGIT regulates antitumor and antiviral CD8(+) T cell effector function. Cancer Cell 26, 923–937. 10.1016/j.ccell.2014.10.018.

7. Schmitz, J.E., Kuroda, M.J., Santra, S., Sasseville, V.G., Simon, M.A., Lifton, M.A., Racz, P., Tenner-Racz, K., Dalesandro, M., Scallon, B.J., et al. (1999). Control of viremia in simian immunodeficiency virus infection by CD8+ lymphocytes. Science 283, 857–860. 10.1126/science.283.5403.857.

8. Belk, J.A., Daniel, B., and Satpathy, A.T. (2022). Epigenetic regulation of T cell exhaustion. Nat Immunol 23, 848–860. 10.1038/s41590-022-01224-z.

9. Zebley, C.C., Gottschalk, S., and Youngblood, B. (2020). Rewriting History: Epigenetic Reprogramming of CD8(+) T Cell Differentiation to Enhance Immunotherapy. Trends Immunol 41, 665–675. 10.1016/j.it.2020.06.008.

10. Chung, H.K., McDonald, B., and Kaech, S.M. (2021). The architectural design of CD8+ T cell responses in acute and chronic infection: Parallel structures with divergent fates. J Exp Med 218. 10.1084/jem.20201730.

11. Sandu, I., and Oxenius, A. (2023). T-cell heterogeneity, progenitor-progeny relationships, and function during latent and chronic viral infections. Immunol Rev 316, 136–159. 10.1111/imr.13203.

12. Zander, R., and Cui, W. (2023). Exhausted CD8(+) T cells face a developmental fork in the road. Trends Immunol 44, 276–286. 10.1016/j.it.2023.02.006.

13. He, R., Hou, S., Liu, C., Zhang, A., Bai, Q., Han, M., Yang, Y., Wei, G., Shen, T., Yang, X., et al. (2016). Follicular CXCR5-expressing CD8(+) T cells curtail chronic viral infection. Nature 537, 412–428. 10.1038/nature19317.

14. Im, S.J., Hashimoto, M., Gerner, M.Y., Lee, J., Kissick, H.T., Burger, M.C., Shan, Q., Hale, J.S., Lee, J., Nasti, T.H., et al. (2016). Defining CD8+ T cells that provide the proliferative burst after PD-1 therapy. Nature 537, 417–421. 10.1038/nature19330.

15. Utzschneider, D.T., Charmoy, M., Chennupati, V., Pousse, L., Ferreira, D.P., Calderon-Copete, S., Danilo, M., Alfei, F., Hofmann, M., Wieland, D., et al. (2016). T Cell Factor 1-Expressing Memory-like CD8(+) T Cells Sustain the Immune Response to Chronic Viral Infections. Immunity 45, 415–427. 10.1016/j.immuni.2016.07.021.

16. Wu, T., Ji, Y., Moseman, E.A., Xu, H.C., Manglani, M., Kirby, M., Anderson, S.M., Handon, R., Kenyon, E., Elkahloun, A., et al. (2016). The TCF1-Bcl6 axis counteracts type I interferon to repress exhaustion and maintain T cell stemness. Sci Immunol 1. 10.1126/sciimmunol.aai8593.

17. Kurtulus, S., Madi, A., Escobar, G., Klapholz, M., Nyman, J., Christian, E., Pawlak, M., Dionne, D., Xia, J., Rozenblatt-Rosen, O., et al. (2019). Checkpoint Blockade Immunotherapy Induces Dynamic Changes in PD-1(-)CD8(+) Tumor-Infiltrating T Cells. Immunity 50, 181–194 e186. 10.1016/j.immuni.2018.11.014.

18. Miller, B.C., Sen, D.R., Al Abosy, R., Bi, K., Virkud, Y.V., LaFleur, M.W., Yates, K.B., Lako, A., Felt, K., Naik, G.S., et al. (2019). Subsets of exhausted CD8(+) T cells differentially mediate tumor control and respond to checkpoint blockade. Nat Immunol 20, 326–336. 10.1038/s41590-019-0312-6.

19. Sade-Feldman, M., Yizhak, K., Bjorgaard, S.L., Ray, J.P., de Boer, C.G., Jenkins, R.W., Lieb, D.J., Chen, J.H., Frederick, D.T., Barzily-Rokni, M., et al. (2018). Defining T Cell States Associated with Response to Checkpoint Immunotherapy in Melanoma. Cell 175, 998–1013 e1020. 10.1016/j.cell.2018.10.038.

20. Beltra, J.C., Manne, S., Abdel-Hakeem, M.S., Kurachi, M., Giles, J.R., Chen, Z., Casella, V., Ngiow, S.F., Khan, O., Huang, Y.J., et al. (2020). Developmental Relationships of Four Exhausted CD8(+) T Cell Subsets Reveals Underlying Transcriptional and Epigenetic Landscape Control Mechanisms. Immunity 52, 825–841 e828. 10.1016/j.immuni.2020.04.014.

21. Hudson, W.H., Gensheimer, J., Hashimoto, M., Wieland, A., Valanparambil, R.M., Li, P., Lin, J.X., Konieczny, B.T., Im, S.J., Freeman, G.J., et al. (2019). Proliferating Transitory T Cells with an Effector-like Transcriptional Signature Emerge from PD-1(+) Stem-like CD8(+) T Cells during Chronic Infection. Immunity 51, 1043–1058 e1044. 10.1016/j.immuni.2019.11.002.

22. Raju, S., Xia, Y., Daniel, B., Yost, K.E., Bradshaw, E., Tonc, E., Verbaro, D.J., Kometani, K., Yokoyama, W.M., Kurosaki, T., et al. (2021). Identification of a T-bet(hi) Quiescent Exhausted CD8 T Cell Subpopulation That Can Differentiate into TIM3(+)CX3CR1(+) Effectors and Memory-like Cells. J Immunol 206, 2924–2936. 10.4049/jimmunol.2001348.

23. Zander, R., Schauder, D., Xin, G., Nguyen, C., Wu, X., Zajac, A., and Cui, W. (2019). CD4(+) T Cell Help Is Required for the Formation of a Cytolytic CD8(+) T Cell Subset that Protects against Chronic Infection and Cancer. Immunity 51, 1028–1042 e1024. 10.1016/j.immuni.2019.10.009.

24. Daniel, B., Yost, K.E., Hsiung, S., Sandor, K., Xia, Y., Qi, Y., Hiam-Galvez, K.J., Black, M., C, J.R., Shi, Q., et al. (2022). Divergent clonal differentiation trajectories of T cell exhaustion. Nat Immunol 23, 1614–1627. 10.1038/s41590-022-01337-5.

25. Giles, J.R., Ngiow, S.F., Manne, S., Baxter, A.E., Khan, O., Wang, P., Staupe, R., Abdel-Hakeem, M.S., Huang, H., Mathew, D., et al. (2022). Shared and distinct biological circuits in effector, memory and exhausted CD8(+) T cells revealed by temporal single-cell transcriptomics and epigenetics. Nat Immunol 23, 1600–1613. 10.1038/s41590-022-01338-4.

26. Kasmani, M.Y., Zander, R., Chung, H.K., Chen, Y., Khatun, A., Damo, M., Topchyan, P., Johnson, K.E., Levashova, D., Burns, R., et al. (2023). Clonal lineage tracing reveals mechanisms skewing CD8+ T cell fate decisions in chronic infection. J Exp Med 220. 10.1084/jem.20220679.

27. Yamauchi, T., Hoki, T., Oba, T., Jain, V., Chen, H., Attwood, K., Battaglia, S., George, S., Chatta, G., Puzanov, I., et al. (2021). T-cell CX3CR1 expression as a dynamic blood-based biomarker of response to immune checkpoint inhibitors. Nat Commun 12, 1402. 10.1038/s41467-021-21619-0.

28. Yan, Y., Cao, S., Liu, X., Harrington, S.M., Bindeman, W.E., Adjei, A.A., Jang, J.S., Jen, J., Li, Y., Chanana, P., et al. (2018). CX3CR1 identifies PD-1 therapy-responsive CD8+ T cells that withstand chemotherapy during cancer chemoimmunotherapy. JCI Insight 3. 10.1172/jci.insight.97828.

29. Kanev, K., and Zehn, D. (2021). Origin and fine-tuning of effector CD8 T cell subpopulations in chronic infection. Curr Opin Virol 46, 27–35. 10.1016/j.coviro.2020.10.003.

30. Grausenburger, R., Bilic, I., Boucheron, N., Zupkovitz, G., El-Housseiny, L., Tschismarov, R., Zhang, Y., Rembold, M., Gaisberger, M., Hartl, A., et al. (2010). Conditional deletion of histone deacetylase 1 in T cells leads to enhanced airway inflammation and increased Th2 cytokine production. J Immunol 185, 3489–3497. 10.4049/jimmunol.0903610.

31. Baazim, H., Schweiger, M., Moschinger, M., Xu, H., Scherer, T., Popa, A., Gallage, S., Ali, A., Khamina, K., Kosack, L., et al. (2019). CD8(+) T cells induce cachexia during chronic viral infection. Nat Immunol 20, 701–710. 10.1038/s41590-019-0397-y.

32. Doherty, P.C., Hou, S., and Southern, P.J. (1993). Lymphocytic choriomeningitis virus induces a chronic wasting disease in mice lacking class I major histocompatibility complex glycoproteins. J Neuroimmunol 46, 11–17. 10.1016/0165-5728(93)90228-q.

33. Moskophidis, D., Lechner, F., Pircher, H., and Zinkernagel, R.M. (1993). Virus persistence in acutely infected immunocompetent mice by exhaustion of antiviral cytotoxic effector T cells. Nature 362, 758–761. 10.1038/362758a0.

34. Yao, C., Sun, H.W., Lacey, N.E., Ji, Y., Moseman, E.A., Shih, H.Y., Heuston, E.F., Kirby, M., Anderson, S., Cheng, J., et al. (2019). Single-cell RNA-seq reveals TOX as a key regulator of CD8(+) T cell persistence in chronic infection. Nat Immunol 20, 890–901. 10.1038/s41590-019-0403-4.

35. Utzschneider, D.T., Gabriel, S.S., Chisanga, D., Gloury, R., Gubser, P.M., Vasanthakumar, A., Shi, W., and Kallies, A. (2020). Early precursor T cells establish and propagate T cell exhaustion in chronic infection. Nat Immunol 21, 1256–1266. 10.1038/s41590-020-0760-z.

36. Alfei, F., Kanev, K., Hofmann, M., Wu, M., Ghoneim, H.E., Roelli, P., Utzschneider, D.T., von Hoesslin, M., Cullen, J.G., Fan, Y., et al. (2019). TOX reinforces the phenotype and longevity of exhausted T cells in chronic viral infection. Nature 571, 265–269. 10.1038/s41586-019-1326-9.

37. Khan, O., Giles, J.R., McDonald, S., Manne, S., Ngiow, S.F., Patel, K.P., Werner, M.T., Huang, A.C., Alexander, K.A., Wu, J.E., et al. (2019). TOX transcriptionally and epigenetically programs CD8(+) T cell exhaustion. Nature 571, 211–218. 10.1038/s41586-019-1325-x.

38. Scott, A.C., Dundar, F., Zumbo, P., Chandran, S.S., Klebanoff, C.A., Shakiba, M., Trivedi, P., Menocal, L., Appleby, H., Camara, S., et al. (2019). TOX is a critical regulator of tumour-specific T cell differentiation. Nature 571, 270–274. 10.1038/s41586-019-1324-y.

39. Seo, H., Chen, J., Gonzalez-Avalos, E., Samaniego-Castruita, D., Das, A., Wang, Y.H., Lopez-Moyado, I.F., Georges, R.O., Zhang, W., Onodera, A., et al. (2019). TOX and TOX2 transcription factors cooperate with NR4A transcription factors to impose CD8(+) T cell exhaustion. Proc Natl Acad Sci U S A 116, 12410–12415. 10.1073/pnas.1905675116.

40. McLane, L.M., Ngiow, S.F., Chen, Z., Attanasio, J., Manne, S., Ruthel, G., Wu, J.E., Staupe, R.P., Xu, W., Amaravadi, R.K., et al. (2021). Role of nuclear localization in the regulation and function of T-bet and Eomes in exhausted CD8 T cells. Cell Rep 35, 109120. 10.1016/j.celrep.2021.109120.

41. Zehn, D., Thimme, R., Lugli, E., de Almeida, G.P., and Oxenius, A. (2022). ’Stem-like’ precursors are the fount to sustain persistent CD8(+) T cell responses. Nat Immunol 23, 836–847. 10.1038/s41590-022-01219-w.

42. Chen, Z., Ji, Z., Ngiow, S.F., Manne, S., Cai, Z., Huang, A.C., Johnson, J., Staupe, R.P., Bengsch, B., Xu, C., et al. (2019). TCF-1-Centered Transcriptional Network Drives an Effector versus Exhausted CD8 T Cell-Fate Decision. Immunity 51, 840–855 e845. 10.1016/j.immuni.2019.09.013.

43. Martin, M.D., and Badovinac, V.P. (2018). Defining Memory CD8 T Cell. Front Immunol 9, 2692. 10.3389/fimmu.2018.02692.

44. Lee, P.P., Fitzpatrick, D.R., Beard, C., Jessup, H.K., Lehar, S., Makar, K.W., Perez-Melgosa, M., Sweetser, M.T., Schlissel, M.S., Nguyen, S., et al. (2001). A critical role for Dnmt1 and DNA methylation in T cell development, function, and survival. Immunity 15, 763–774. 10.1016/s1074-7613(01)00227-8.

45. Pircher, H., Burki, K., Lang, R., Hengartner, H., and Zinkernagel, R.M. (1989). Tolerance induction in double specific T-cell receptor transgenic mice varies with antigen. Nature 342, 559–561. 10.1038/342559a0.

46. Srinivas, S., Watanabe, T., Lin, C.S., William, C.M., Tanabe, Y., Jessell, T.M., and Costantini, F. (2001). Cre reporter strains produced by targeted insertion of EYFP and ECFP into the ROSA26 locus. BMC Dev Biol 1, 4. 10.1186/1471-213x-1-4.

47. Hameyer, D., Loonstra, A., Eshkind, L., Schmitt, S., Antunes, C., Groen, A., Bindels, E., Jonkers, J., Krimpenfort, P., Meuwissen, R., et al. (2007). Toxicity of ligand-dependent Cre recombinases and generation of a conditional Cre deleter mouse allowing mosaic recombination in peripheral tissues. Physiol Genomics 31, 32–41. 10.1152/physiolgenomics.00019.2007.

48. Chen, Y., Zander, R.A., Wu, X., Schauder, D.M., Kasmani, M.Y., Shen, J., Zheng, S., Burns, R., Taparowsky, E.J., and Cui, W. (2021). BATF regulates progenitor to cytolytic effector CD8(+) T cell transition during chronic viral infection. Nat Immunol 22, 996–1007. 10.1038/s41590-021-00965-7.

49. Wherry, E.J., Ha, S.J., Kaech, S.M., Haining, W.N., Sarkar, S., Kalia, V., Subramaniam, S., Blattman, J.N., Barber, D.L., and Ahmed, R. (2007). Molecular signature of CD8+ T cell exhaustion during chronic viral infection. Immunity 27, 670–684. 10.1016/j.immuni.2007.09.006.

50. Seto, E., and Yoshida, M. (2014). Erasers of histone acetylation: the histone deacetylase enzymes. Cold Spring Harb Perspect Biol 6, a018713. 10.1101/cshperspect.a018713.

51. Yang, X.J., and Seto, E. (2007). HATs and HDACs: from structure, function and regulation to novel strategies for therapy and prevention. Oncogene 26, 5310–5318. 10.1038/sj.onc.1210599.

52. Choudhary, C., Kumar, C., Gnad, F., Nielsen, M.L., Rehman, M., Walther, T.C., Olsen, J.V., and Mann, M. (2009). Lysine acetylation targets protein complexes and co-regulates major cellular functions. Science 325, 834–840. 10.1126/science.1175371.

53. Ellmeier, W., and Seiser, C. (2018). Histone deacetylase function in CD4(+) T cells. Nat Rev Immunol 18, 617–634. 10.1038/s41577-018-0037-z.

54. Baxter, A.E., Huang, H., Giles, J.R., Chen, Z., Wu, J.E., Drury, S., Dalton, K., Park, S.L., Torres, L., Simone, B.W., et al. (2023). The SWI/SNF chromatin remodeling complexes BAF and PBAF differentially regulate epigenetic transitions in exhausted CD8(+) T cells. Immunity 56, 1320–1340 e1310. 10.1016/j.immuni.2023.05.008.

55. Beltra, J.C., Abdel-Hakeem, M.S., Manne, S., Zhang, Z., Huang, H., Kurachi, M., Su, L., Picton, L., Ngiow, S.F., Muroyama, Y., et al. (2023). Stat5 opposes the transcription factor Tox and rewires exhausted CD8(+) T cells toward durable effector-like states during chronic antigen exposure. Immunity 56, 2699–2718 e2611. 10.1016/j.immuni.2023.11.005.

56. Barber, D.L., Wherry, E.J., Masopust, D., Zhu, B., Allison, J.P., Sharpe, A.H., Freeman, G.J., and Ahmed, R. (2006). Restoring function in exhausted CD8 T cells during chronic viral infection. Nature 439, 682–687. 10.1038/nature04444.

57. Tsui, C., Kretschmer, L., Rapelius, S., Gabriel, S.S., Chisanga, D., Knopper, K., Utzschneider, D.T., Nussing, S., Liao, Y., Mason, T., et al. (2022). MYB orchestrates T cell exhaustion and response to checkpoint inhibition. Nature 609, 354–360. 10.1038/s41586-022-05105-1.

58. Moser, M.A., Hagelkruys, A., and Seiser, C. (2014). Transcription and beyond: the role of mammalian class I lysine deacetylases. Chromosoma 123, 67–78. 10.1007/s00412-013-0441-x.

59. Bergthaler, A., Flatz, L., Hegazy, A.N., Johnson, S., Horvath, E., Lohning, M., and Pinschewer, D.D. (2010). Viral replicative capacity is the primary determinant of lymphocytic choriomeningitis virus persistence and immunosuppression. Proc Natl Acad Sci U S A 107, 21641–21646. 10.1073/pnas.1011998107.

60. Welsh, R.M., and Seedhom, M.O. (2008). Lymphocytic choriomeningitis virus (LCMV): propagation, quantitation, and storage. Curr Protoc Microbiol Chapter 15, Unit 15A 11. 10.1002/9780471729259.mc15a01s8.

61. Battegay, M., Cooper, S., Althage, A., Banziger, J., Hengartner, H., and Zinkernagel, R.M. (1991). Quantification of lymphocytic choriomeningitis virus with an immunological focus assay in 24- or 96-well plates. J Virol Methods 33, 191–198. 10.1016/0166-0934(91)90018-u.

62. Gulich, A.F., Rica, R., Tizian, C., Viczenczova, C., Khamina, K., Faux, T., Hainberger, D., Penz, T., Bosselut, R., Bock, C., et al. (2021). Complex Interplay Between MAZR and Runx3 Regulates the Generation of Cytotoxic T Lymphocyte and Memory T Cells. Front Immunol 12, 535039. 10.3389/fimmu.2021.535039.

63. Buenrostro, J.D., Giresi, P.G., Zaba, L.C., Chang, H.Y., and Greenleaf, W.J. (2013). Transposition of native chromatin for fast and sensitive epigenomic profiling of open chromatin, DNA-binding proteins and nucleosome position. Nat Methods 10, 1213–1218. 10.1038/nmeth.2688.

64. Corces, M.R., Buenrostro, J.D., Wu, B., Greenside, P.G., Chan, S.M., Koenig, J.L., Snyder, M.P., Pritchard, J.K., Kundaje, A., Greenleaf, W.J., et al. (2016). Lineage-specific and single-cell chromatin accessibility charts human hematopoiesis and leukemia evolution. Nat Genet 48, 1193–1203. 10.1038/ng.3646.

65. Consortium, E.P. (2012). An integrated encyclopedia of DNA elements in the human genome. Nature 489, 57–74. 10.1038/nature11247.

